# Structures of a P4-ATPase lipid flippase in lipid bilayers

**DOI:** 10.1101/2020.03.14.992305

**Authors:** Yilin He, Jinkun Xu, Xiaofei Wu, Long Li

## Abstract

Type 4 P-type ATPases (P4-ATPases) are a group of key enzymes maintaining lipid asymmetry of eukaryotic membranes. Phospholipids are actively and selectively flipped by P4-ATPases from the exoplasmic leaflet to the cytoplasmic leaflet. How lipid flipping is coupled with ATP-hydrolysis by P4-ATPases is poorly understood. Here, we report the electron cryo-microscopy structures of a P4-ATPase, Dnf1-Cdc50 from *Chaetomium thermophilum*, which had been reconstituted into lipid nanodiscs and captured in two transport intermediate states. The structures reveal that transmembrane segment 1 of Dnf1 becomes highly flexible during lipid transport. The local lipid bilayers are distorted to facilitate the entry of the phospholipid substrates from the exoplasmic leaflet to a cross-membrane groove. During transport, the lipid substrates are relayed through four binding sites in the groove which constantly shields the lipid polar heads away from the hydrophobic environment of the membranes.

## Introduction

Phospholipid molecules are unevenly distributed in membrane bilayers of eukaryotic cells ^1,2^. Phosphatidylethanolamine (PE) and phosphatidylserine (PS) are concentrated in the cytoplasmic leaflet, whereas phosphatidylcholine (PC) is enriched in the exoplasmic leaflet (lumenal or extracellular leaflet)^3^. The asymmetric distribution of phospholipids is critical for numerous cellular processes, such as cell signaling, apoptosis, and blood coagulation ^4,5^. Lipid asymmetry is created and maintained by ATP-driven lipid translocases, such as lipid flippases that move specific lipids from the exoplasmic leaflet to the cytoplasmic leaflet, and lipid floppases which move lipids from the cytoplasmic to the exoplasmic leaflet. Lipid asymmetry can be disrupted by lipid scramblases which, upon activation, facilitate the rapid bi-directional movement of lipids across the two leaflets. Extensive studies in the past few years revealed that type 4 P-type ATPases (P4-ATPases) are the phospholipid flippases, a group of ATP-binding cassette (ABC) transporters are the floppases, and transmembrane protein 16F (TMEM16F) could function as the non-specific lipid scramblase ^6^. Among the lipid translocases, P4-ATPases belong to the P-type ATPase family. Most members in the family transport small metal ions across membranes, whereas P4-ATPases specifically transport phospholipids that are at least ten times larger than the metal ions ^7^. Therefore, it was postulated that P4-ATPases had a distinct mechanism from other P-type ATPases for substrate transport ^8-11^.

P4-ATPases are conserved in eukaryotes. *Saccharomyces cerevisiae* (*S. cerevisiae*) has five P4-ATPases characterized, namely Drs2, Dnf1, Dnf2, Dnf3, and Neo1, whereas 14 P4-ATPases are identified in human^12,13^. Studies on mouse models showed that deficiencies in P4-ATPases resulted in neurological disorders, liver diseases, immunological problems, and diabetes ^14^. On the cellular level, P4-ATPases are required in vesicular traffickings, such as the formation of endosomes and post-Golgi secretory vesicles ^15^. Each P4-ATPase has its preferred lipid substrates. For example, yeast Drs2 translocates PS and PE, whereas Dnf1 and Dnf2 prefer PC, PE, and glucosylceramide ^16^. Most P4-ATPases form heterodimers with a β-subunit from the CDC50 family. The β-subunit is required for proper folding, sub-cellular targeting, and lipid flipping of P4-ATPases ^17,18^.

Lipid flipping by P4-ATPases is coupled with ATP hydrolysis and enzyme phosphorylation/dephosphorylation. Similar to other P-type ATPases, such as the sarcoplasmic reticulum calcium pump (SERCA) and Na/K-ATPases, P4-ATPases undergo the E1-E2 state transition during lipid flipping cycles ^19^. Lipid substrate binding is coupled with dephosphorylation in the low energy E2P state whereas the role of the E1 state in lipid flipping is less clear. The timing when the phospholipids are picked up from the exoplasmic leaflet and released to the cytoplasmic leaflet during the E1-E2 transition is obscure. The lipid flipping pathway in P4-ATPases is also under debate. A “two-gate” model suggests that specific phospholipids are recognized at the entry and exit gates by the residues clustering at the exoplasmic and cytoplasmic ends of transmembrane segments (TMs) 1-4. The polar heads of the lipid substrates could slide through a groove formed by TMs 1, 3, and 4 during flipping ^20,21^. A “hydrophobic gate” model proposed a cross-membrane groove bordered by TMs 1, 2, 4, and 6. A highly conserved isoleucine residue (I364 in bovine ATP8A2) and a few hydrophobic residues nearby act as a hydrophobic gate to control lipid movements ^22^. A third model proposed that a central cavity between TMs 3, 5, and 6 could accommodate the head groups of phospholipids during transporting across the membranes^23^.

Recently, the electron cryo-microscopy (cryo-EM) structures of *S. cerevisiae* Drs2-Cdc50 (scDrs2-Cdc50)^24,25^ and human ATP8A1-CDC50a (hATP8A1-CDC50a)^26^ solubilized in detergents were reported. The structures of scDrs2-Cdc50 were determined in the E2P state and the hATP8A1-CDC50a structures in several E1 and E2 intermediate states. However, the subtle movements of TMs among the structures make it hard to draw a consensus model for lipid flipping. Here, we report the cryo-EM structures of a P4-ATPase from a thermophilic fungus reconstituted into lipid nanodiscs. The structures are determined in the presence of β, γ-methyleneadenosine 5′-triphosphate (AMPPCP) and beryllium fluoride (BeF_3_^−^), representing the E1-ATP and E2P states, respectively. The large conformational changes between the two states suggest a mechanism underlying lipid flipping by P4-ATPases during the E1-E2 transition.

## Results

### Overall structures

P4-ATPases from a thermophilic fungus, *Chaetomium thermophilum* (*C. thermophilum*) were cloned for structural studies. Protein BLAST search identified three P4-ATPases, including one *S. cerevisiae* Drs2 homolog (ctDrs2), one Dnf1 and Dnf2 homolog (ctDnf1), and one Dnf3 homolog (ctDnf3) (Fig. S1). Only one CDC50 protein was found in the *C. thermophilum* genome (ctCdc50). After initial screening, ctDnf1 was chosen to co-express with ctCdc50 in yeast. The purified ctDnf1-Cdc50 heterodimer showed phospholipid-dependent ATPase activity (Fig. S2). Both phosphatidylcholine (PC) and phosphatidylserine (PS) could stimulate ATPase activity, with PC being slightly better than PS at high concentrations (Fig. S2d). In contrast, scDrs2 was shown to be selectively stimulated by PS^27,28^. Therefore, ctDnf1 is likely to have different substrate preference from scDrs2, maybe preferring PC to PS, similar to scDnf1 and scDnf2^20,21,29^. To mimic the native lipid membrane environment, ctDnf1-Cdc50 was reconstituted into lipid nanodiscs^30^. The complex was supplemented with AMPPCP (E1-ATP) or BeF_3_^−^ (E2P) and subject to cryo-EM single-particle analyses. The structures were determined to a resolution of 3.5 Å for E1-ATP and of 3.4 Å for E2P (Fig. S3-5).

Similar to scDrs2 and hATP8A1, ctDnf1 has all the typical domains of P-type ATPases, namely the actuator domain (A), the nucleotide-binding domain (N), the phosphorylation domain (P), and the membrane domain (M) (Fig. 1). In both E1-ATP and E2P structures, ctCdc50 has extensive interactions with ctDnf1. On the cytoplasmic side, the N-terminal peptide (residues 23-46) of ctCdc50 runs along one face of ctDnf1, interacting with the cytosolic loops of ctDnf1, including the segment connecting TM4 and the P domain, the loop between TMs 6 and 7 (L6/7), L8/9, and a C-terminal amphipathic helix that was suggested to undergo conformational changes upon PI4P activation in scDrs2^24,25^ (Fig. S6). The C-terminal tail of ctCdc50 (residues 384-398), which is invisible in the scCdc50 and hCDC50a structures, could be traced to run towards ctDnf1. Thus, the C-terminal tail and the N-terminal peptide of ctCdc50 are in such a conformation that sandwiches the C-terminal amphipathic helix of ctDnf1 (Fig. S6). The two TMs of ctCdc50 form hydrophobic interactions with TM7 and TM10 of ctDnf1. The ectodomain of ctCdc50 has the largest interfaces with ctDnf1, interacting with all the exoplasmic loops except L1/2 (Fig. 1c). Thus, ctCdc50 acts as a 3-way clamp to hold TMs 3-10 of ctDnf1 in a relatively fixed conformation in both E1-ATP and E2P structures.

**Fig. 1.**
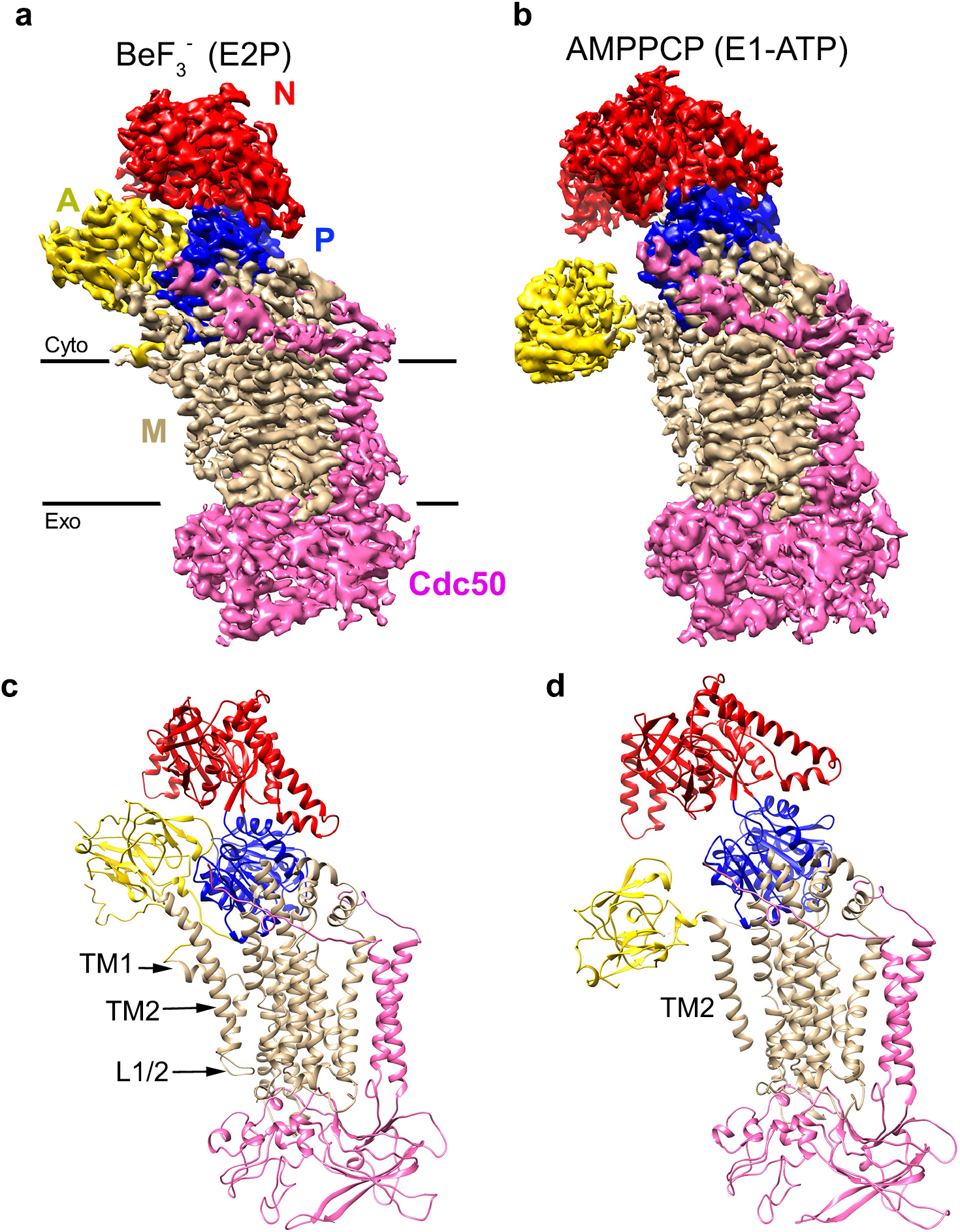
Structures of ctDnf1-Cdc50. **a**, Cryo-EM map of ctDnf1-Cdc50 with BeF_3_^-^. The A, N, P, and M domains and Cdc50 are labeled and colored yellow, red, blue, tan, and pink, respectively. **b**, Cryo-EM map of ctDnf1-Cdc50 with AMPPCP. Colors are the same as in **a. c**, Ribbon diagram of ctDnf1-Cdc50 with BeF_3_^-^. TM1, TM2, and L1/2 are labeled. **d**, Ribbon diagram of ctDnf1-Cdc50 with AMPPCP. TM2 is labeled.

### Conformational changes of ctDnf1 between the E1-ATP and E2P states

Comparison of the ctDnf1-Cdc50 structures in the E1-ATP and the E2P states shows that ctCdc50 and TMs 3-10 of ctDnf1 are quite similar (root mean square deviation = 0.79 Å), whereas the largest differences are seen in the N domain, the A domain, and TMs 1-2 of ctDnf1 (Fig. 2). In the E2P structure, BeF_3_^−^ is bound at the phosphorylation site D606 (Fig. S5h), same as in the typical E2P structures of P-type ATPases^24,26,31^. The A domain associates tightly with the N and P domains. Both A and N domains are in an upright conformation (Fig. 1a and 1c). In the E1-ATP structure, AMPPCP adapts an extended conformation (Fig. S5i), different from the bent conformations observed in hTAP8A1 and other P-type ATPases (Fig. S7)^26,31^. The extended conformation suggests the structure may represent a distinct active E1-ATP state not observed before^32,33^. As a result, the N domain rotates by 35 ° compared to its orientation in the E2P structure (Fig. 2c). The A domain has even larger conformational changes. It moves close to the M domain, no longer associated with the N and P domains (Fig. 1b and 1d). The displacement of A domain is about 30 Å (Fig. 2b), the largest among the known structural changes of P-type ATPases during the E2-E1 transition. TMs 1 and 2 that are directly connected to the A domain have high flexibility in the E1-ATP state. The entire TM1 including the amphipathic helix (residues 122-134) and L1/2 are invisible in the density map, indicating its high flexibility. The C-terminal segment of TM2 that extend to domain A (residues 185-192) is bent and largely disordered. The TM2 density in E1-ATP is also weaker than in E2P, suggesting a more flexible TM2 in E1-ATP. Indeed, in the refined models, the average B factors of TM2 (residues 158-184) are 38.5 in E2P and 71.7 in E1-ATP.

**Fig. 2.**
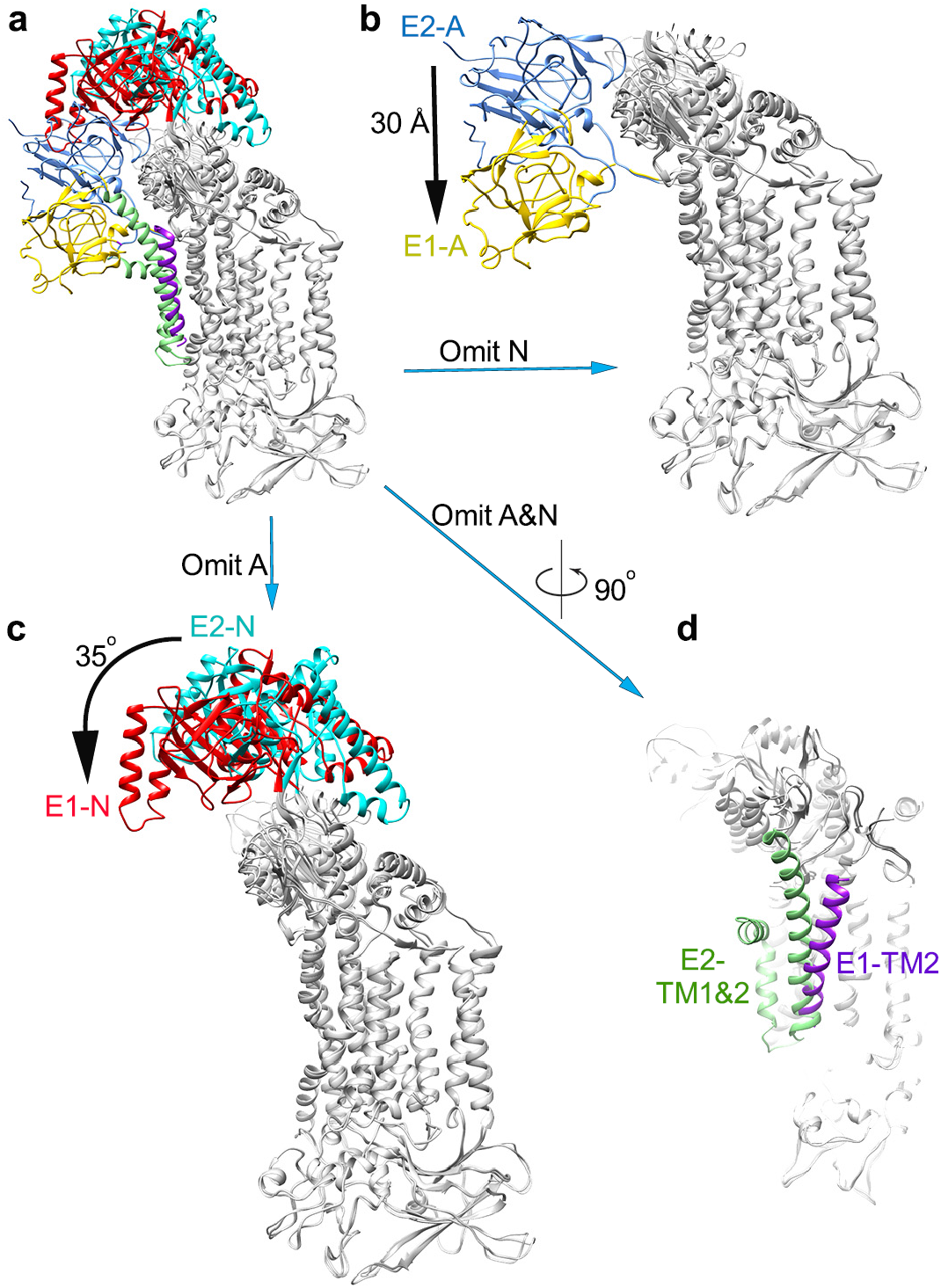
Comparison of ctDnf1-Cdc50 structures in the E1-ATP and E2P states. **a**, Overlay of the E1-ATP and E2P structures by superimposing TMs 3-10 of ctDnf1 and ctCdc50 (grey). The A domain is yellow in E1-ATP and blue in E2P. The N domain is red in E1-ATP and cyan in E2P. TM2 is purple in E1-ATP and TMs 1 and 2 are green in E2P. **b**, same as **a**, except the N domains are omitted for clarity. The motion distance of the A domain between the E1-ATP an E2P states is labeled. **c**, same as **a**, except the A domains are omitted. The movement of the N domain between the E1 and E2 states is indicated. **d**, same as **a**, except rotating by 90 degrees and the A and N domains are omitted to show the movements of TMs 1 and 2 between the E1-ATP and E2P states.

### Membrane distortion by ctDnf1-Cdc50

The structures of ctDnf1-Cdc50 in lipid nanodiscs not only allow us to visualize conformational changes of the flippase during the E1-E2 transition, but also to analyze the structural changes of the membranes. The cryo-EM density maps clearly show that the M domain is wrapped in lipid nanodiscs (Fig. 3). In the E2P structure (Fig. 3a-c), the amphipathic helix of TM1 lies on the surface of the lipid membranes, extending to the brink of the nanodisc (Fig. 3c). The side wall of the nanodisc close to TMs 1 and 2 is not well sealed, indicating that the local lipid bilayers might be disturbed (Fig. 3b). Nevertheless, the nanodisc has a roughly even thickness of ∼36 Å.

**Fig. 3.**
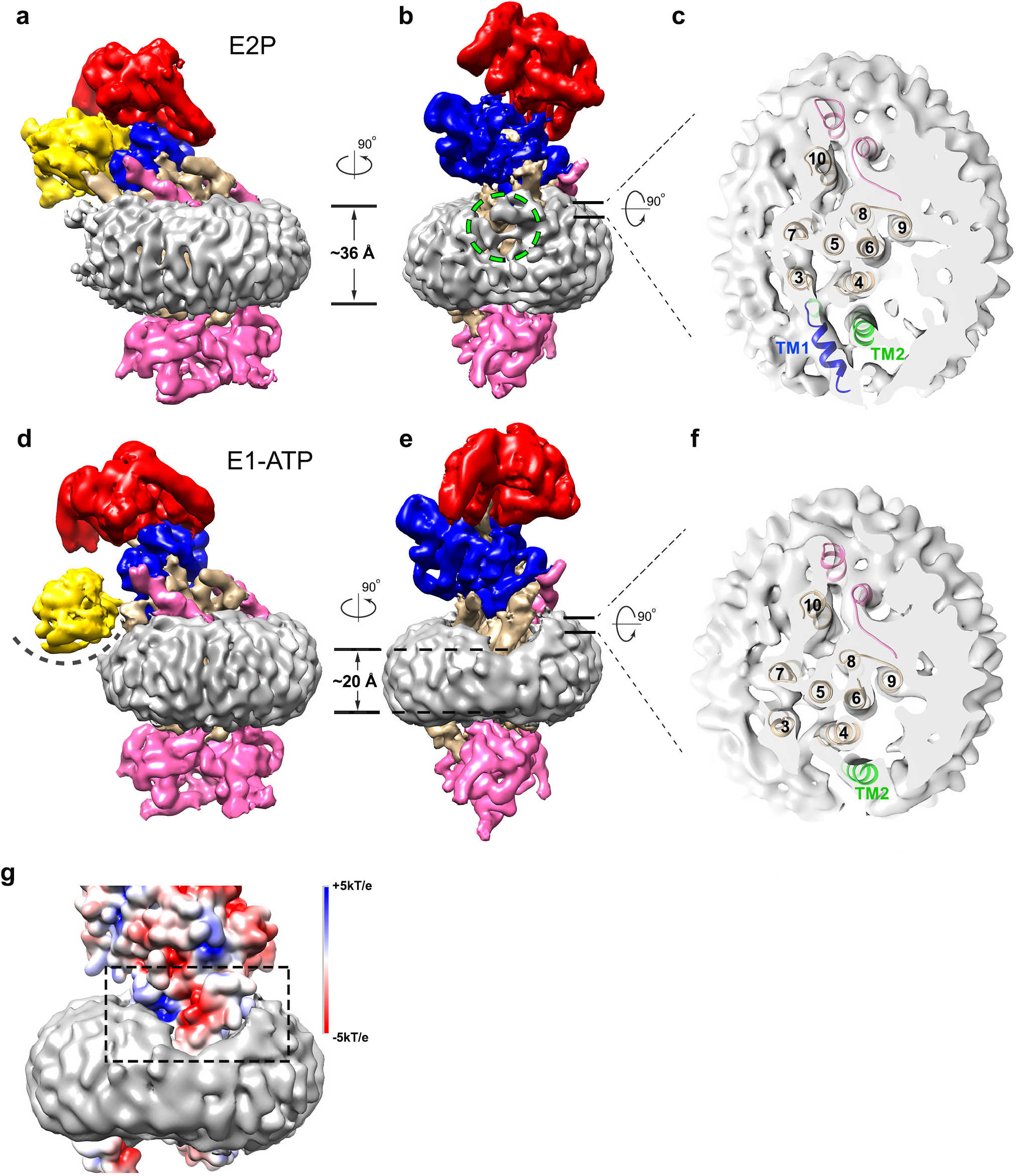
Membrane structures in the E1-ATP and E2P state. **a**, Cryo-EM density map of the E2P state with the lipid nanodisc. The map is low pass filtered to 6 Å. Colors are as in Fig. 1, with the addition of lipid nanodisc density (grey). **b**, 90-degree rotation of **a**, showing the nanodisc density around TMs 1 and 2 (green dashed circle). The A domain is omitted for clarity. **c**, Cut-away top view of the nanodisc. TMs are labeled. **d**, Cryo-EM density map of the E1-ATP state with the lipid nanodisc. The curved dashed line indicates the depression that is required to fit the A domain on the surface of the membranes. **e**, 90-degree rotation of **d**, showing the local membrane thinning. **f**, Cut-away top view of the nanodisc. **g**, Zoom-in view of **e**. The surface of ctDnf1 is colored according to the electrostatic potential. The black dashed frame highlights the distorted membranes and the negatively charged patch.

In the E1-ATP structure (Fig. 3d-f), the lipid bilayers close to TMs 1 and 2 are dramatically distorted. The local membranes shrink to ∼20 Å in thickness (Fig. 3e). Lipid thinning exposes a negatively charged segment of TM2 (residues 174-181) that is shielded away from the lipids in the E2P structure by interacting with the amphipathic helix of TM1. In the context of the native membranes, lipid thinning would result in a depression on the membrane surface to fit the A domain (Fig. 3d). The structure suggests that the large displacement of the A domain in the E1-ATP state might dislodge TM1, which becomes highly flexible and expose a negatively charged patch (Fig. 3g). The patch forming residues, D175, E178 of TM1 and E561 of TM4 are highly conserved among P4-ATPases (Fig. S1). The flexible TM1 and the charged patch might have the ability to distort the local membranes whose thickness is shrunk by almost a half (Fig. 3g). The A domain on the membrane surface may help to distort the local bilayers as well. Interestingly, local membrane thinning and distortion have also been observed for the lipid scramblase TMEM16F ^34^. It should be noted that membrane thinning and distortion are seen at the edge of the nanodisc where the membrane scaffold protein (MSP) wraps around the lipids. Although the MSP could not be resolved in the density maps, it may be involved during local membrane changes as well. Further studies are required to clarify the issue.

### Lipid recognition

Distinct lipid binding sites are identified in the E2P and E1-ATP structures of ctDnf1-Cdc50. In the E2P structure, a phospholipid molecule is found in a cavity formed by TMs 2, 4, and 6 in the exoplasmic half of the M domain (E2-site1) (Fig. 4a, b, Fig. S8a). A prominent density blob corresponding to a phospholipid snorkels deep into the positively charged cavity. A PC molecule is modeled in the density, though it could be other kinds of phospholipids as the resolution is insufficient to distinguish the head groups. The head group is surrounded by residues Q549 and N550 of TM4 and N1153 of TM6 (Fig. 4a). The following acyl chains bent almost 90°, extending into the membranes. A similar lipid binding site is observed in the E2Pi structure of hATP8A1 (AlF_4_^−^ bound)^26^ (Fig. S8b). In that structure, the serine group of the modeled PS molecule interacts with N352 of hATP8A1 (equivalent to Q549 in ctDnf1), whereas the head group density in our structure points deeper into the cavity (Fig. S8c). In addition, the acyl chains of PS run along hATP8A1 pointing to the cytosolic side, unlike the acyl chains that go into the middle of the lipid bilayers in our structure. Therefore, the E2-site1 is likely for early recognition of the phospholipid substrates that have been picked up by the flippase, but are still located in the exoplasmic leaflet. A second lipid molecule is identified at the cytoplasmic end of the groove formed by TMs 2, 4, and 6 (E2-site2) (Fig. 4a, b). The phosphate head group is at the water-membrane interface, clamped by R181 of TM2, F569 and Y572 of TM4, which are not conserved among P4-ATPase though. The acyl chains run along the groove, with the same orientation as the lipids in the cytoplasmic leaflet. Therefore, it might represent the phospholipid that has finished flipping and be ready to be released to the cytoplasmic leaflet. It remains to be seen whether the site is conserved among P4-ATPases.

**Fig. 4.**
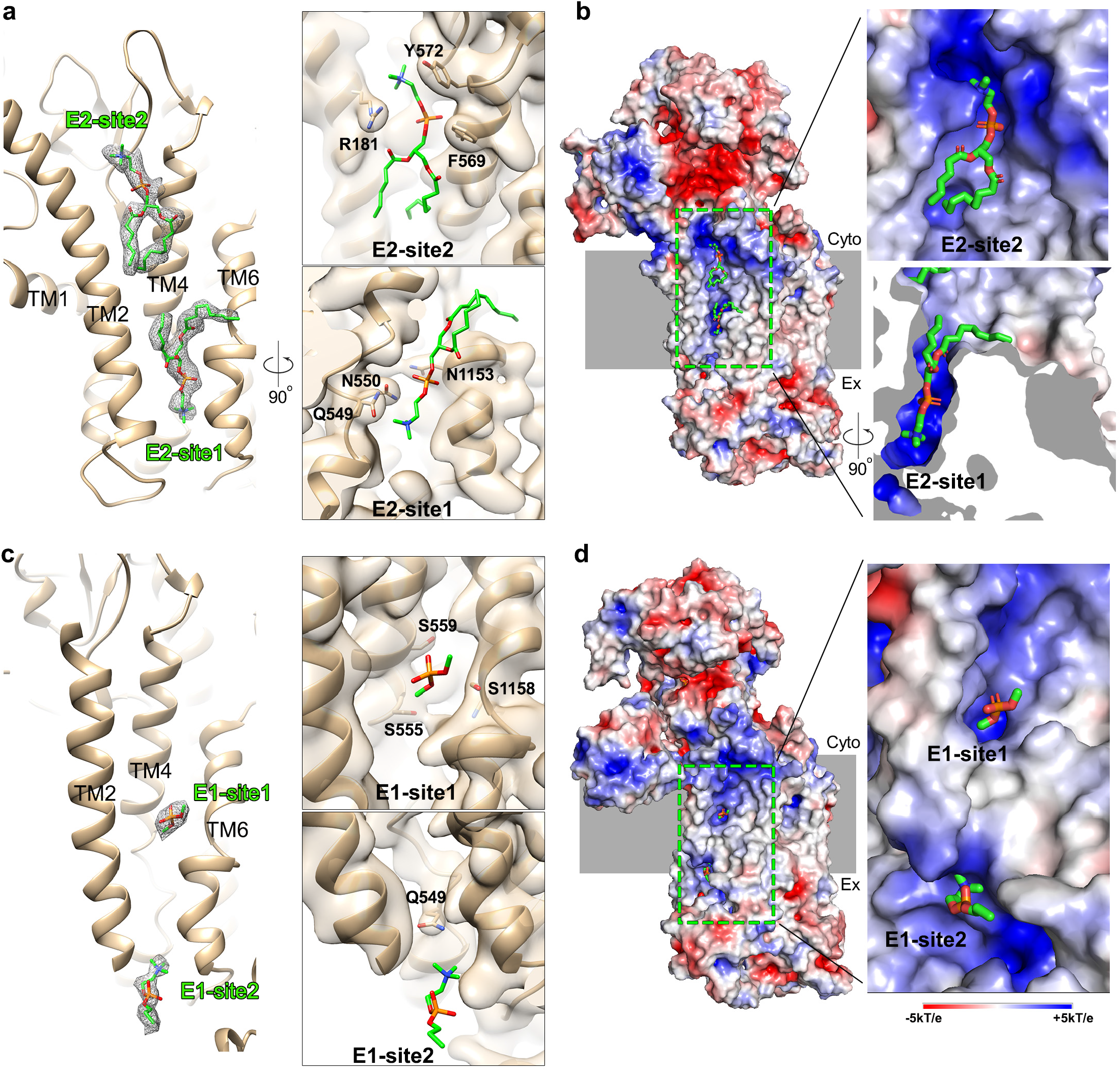
Lipid substrate binding sites. **a**, Lipid binding sites in E2P. ctDnf1 is shown as ribbon. The density of the lipids is shown as grey meshes at 1.5 σ. Two modeled PC and their interacting residues are shown as sticks. The left panel shows the overview and the right panels show the magnified views of each site. TMs, residues, and binding sites are labeled. **b**, Electrostatic potential surfaces of the flippase in E2P, showing the environment of the lipid binding sites. The electrostatic potential surfaces are calculated using APBS with the default setting in PyMOL. The membrane cartoon is colored grey. The left panel shows the overview and the right panels show the magnified views of each site. **c**-**d**, same as **a** and **b**, except showing the binding sites in the E1-ATP structure. A phosphate head group and a lyso-PC are modelled in site 1 and 2, respectively.

In the E1-ATP structure, a relatively small but distinct density blob is found halfway through the membranes in the groove bordered by TMs 2, 4, and 6 (E1-site1) (Fig. 4c, d), two helices away from E2-site1. The density could be the head group of a phospholipid or a hydrophilic molecule as it is close to S1158 of TM6 and points to highly conserved S555 and S559 of TM4 in the groove (Fig. S1). The elongated density blobs are also found at the same locations in the scDrs2 and hATP8A1 structures (Fig. S9), though they are not discussed in the papers^24-26^. This spot may serve as a transient yet conserved binding site in the middle of the phospholipid flipping pathway. A mutation in ATP8B1, S403Y (equivalent to S555 in ctDnf1), was found in the patients with progressive familial intrahepatic cholestasis type 1 (PFIC1)^35^, suggesting an important role of the site in lipid flipping.

The distorted lipid bilayers and the disordered TM1 and L1/2 in the E1-ATP structure pose an opportunity for phospholipids to enter the groove between TMs 2, 4, and 6. Indeed, a rod-like density blob is found at the opening between the exoplasmic ends of TMs 2 and 6, on the surface of the membranes (Fig. 4c, d) (E1-site2). The side chains of Q549 and N550 of TM4 in the groove are in proximity to the density, in a position ready to pick a phospholipid from the membranes. Due to the low local resolution, it is not entirely certain whether the density blob represents a phospholipid that is entering the groove. Nevertheless, the wide opening between TMs 2 and 6 and the local positive charges (Fig. 4d) suggest it is a promising lipid entry site.

The difference in lipid binding between the E1-ATP and E2P structures is mainly caused by the conformational changes of TM2 and TM4. As E2P shifts to E1-ATP, E2-site1 becomes too small to accommodate the head group of a phospholipid (Fig. 5). The local main chain changes of TM4 move the side chains of Q549 and N550 towards the membranes, occupying the space that is filled by the head group of the phospholipid in the E2P structure (Fig. 5b). The shift of TM2 towards TM4 further narrows the path that allows the acyl chains of the lipid to come out from the cavity, while creates a shallow lipid binding site, E1-site1. The disordered L1/2 and the exoplasmic end of TM2 in the E1-ATP structure would allow the phospholipids to enter from E1-site2 (Fig. 5a), where the “cavity-occupying” Q549 and N550 have access to the phospholipids through the opening between TMs 2 and 6 (Fig. 4c, 5a). On the cytoplasmic side, the large shift of TM2 towards TM4 disrupts E2-site2, leaving no space for lipid binding (Fig. 5a).

**Fig. 5.**
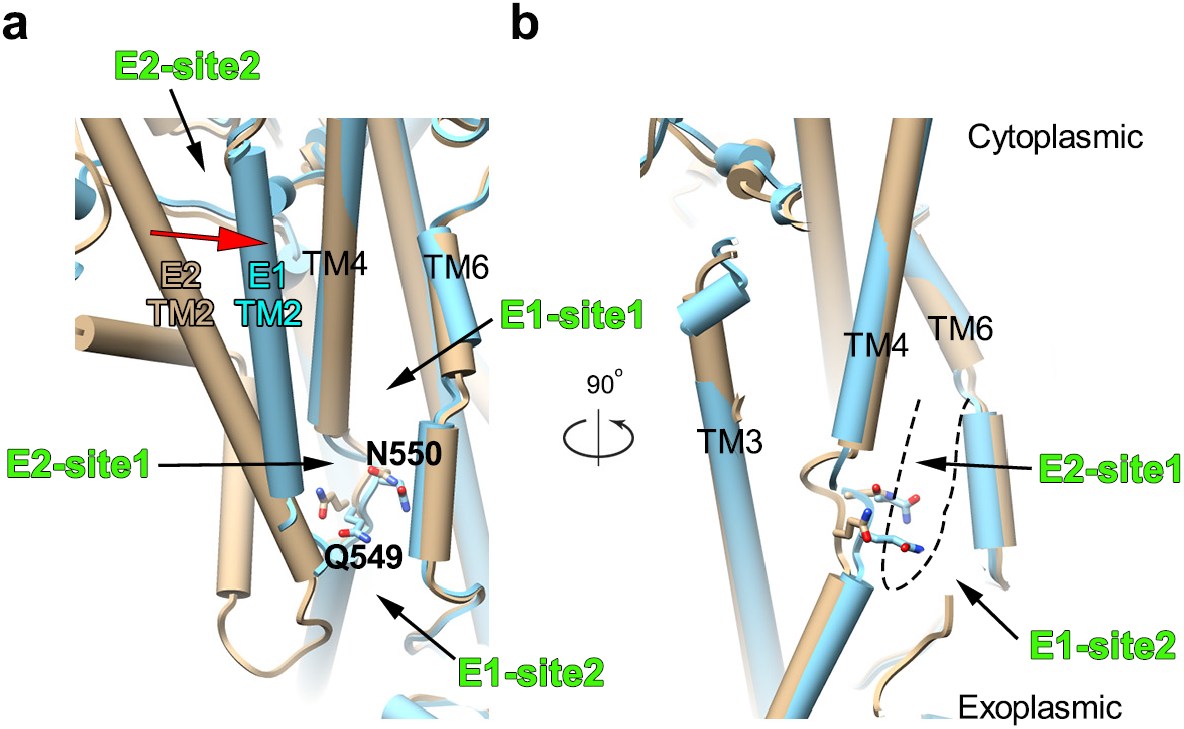
Conformational changes of the lipid substrate binding sites between E1-ATP and E2P. **a**, Superimposition of E1-ATP (cyan cylinders) and E2P (tan cylinders), showing the groove bordered by TMs 2, 4, and 6. Q549 and N550 that interact with the lipid substrate are shown as sticks. The movement of TM2 during the E2-E1 transition is indicated by a red arrow. The lipid binding sites are marked. **b**, Side view of the groove. The dashed line outlines E2-site1 which is disrupted and occupied by Q549 and N550 in E1-ATP.

The E1-ATP and E2P structures also have a lipid-binding site in common in a cavity formed by TMs 7, 8, and 10, on the opposite side of TMs 2, 4, and 6 (Fig. S10). Phospholipid density is observed at the cytoplasmic border of the membranes. The cavity was suggested to be a PI4P binding site for scDrs2 activation^24^. However, in our structures, the head group does not insert as deep as PI4P in scDrs2 E2P^inter^ or E2P^active^. Instead, it is likely to be the head group of PS as modeled in scDrs2 E2P^inhib 24^.

## Discussion

### Phospholipid flipping coupled with the E1-E2 transition

Four lipid-binding sites are identified in the groove formed by TMs 2, 4, and 6, two from E1-ATP and two from E2P. The four sites arranged in such a way that they could relay the phospholipid substrates through the groove during the E1-E2 transition (Fig. 6). Conformational changes of TM1 and TM2 guide the phospholipids to move from the exoplasmic leaflet to the cytoplasmic leaflet. ATP binding to the lipid flippase in the E1-ATP state detaches the A domain from the N and P domains. The large motion of the A domain increases the flexibility of TMs 1 and 2 and exposes a negatively charged patch formed by the residues from TMs 2 and 4. The local lipid bilayers are distorted and the membranes are thinned by almost a half. In the distorted membranes, the phospholipid molecules are more likely to tilt parallel to the membrane plane. The lipid head groups are in an inward-facing orientation, ready to enter the groove via E1-site2 (Fig. 6a). In the E2P state, the A domain is associated with the P and N domains tightly, leading to a relatively rigid conformation of TMs 1 and 2. The local lipid bilayer structures are restored. TMs 2, 4, and 6 create a cavity (E2-site1) in the exoplasmic leaflet for shielding the polar head group of the phospholipid that has been picked up from E1-site2 (Fig. 6b). E2-site1 is disrupted in the E1-ATP state as TM2 moves towards TMs 4 and 6 and two polar residues of TM4 (Q549 and N550) lean towards the membranes. The lipid head group is likely to be squeezed out and to move forward to the cytoplasmic leaflet via E1-site1, a shallow hydrophilic cleft (Fig. 6c). Finally, the phospholipid reaches E2-site2 and is held by a clamp between TMs 2 and 4 in a flipped conformation (Fig. 6d). The phospholipid substrate is ready to be laterally released into the cytoplasmic leaflet when the clamp is disrupted in the E1-ATP state. In the scenario proposed above, it takes two E1-E2 cycles to flip one phospholipid substrate, but costs one ATP molecule per phospholipid on average because two phospholipids are present at the same time in each state. However, as we are missing several intermediate states, e.g. E2 and E1P, the exact cycle number and ATP cost per substrate need further investigation. During lipid transport, the hydrophilic head group of the phospholipid substrate is constantly protected from the hydrophobic environment by sliding through the binding sites in the positively charged groove (Fig. 4b, d). The groove provides the only continuous hydrophilic pathway in the M domain during the E1-E2 transition (Compare Fig. 4b, d and Fig. S10c). The positive charges are mainly contributed by K174, R181, and K1121 on the TMs. The highly conserved K1121 has been shown to be important for substrate binding and ATPase activity^19^.

**Fig. 6.**
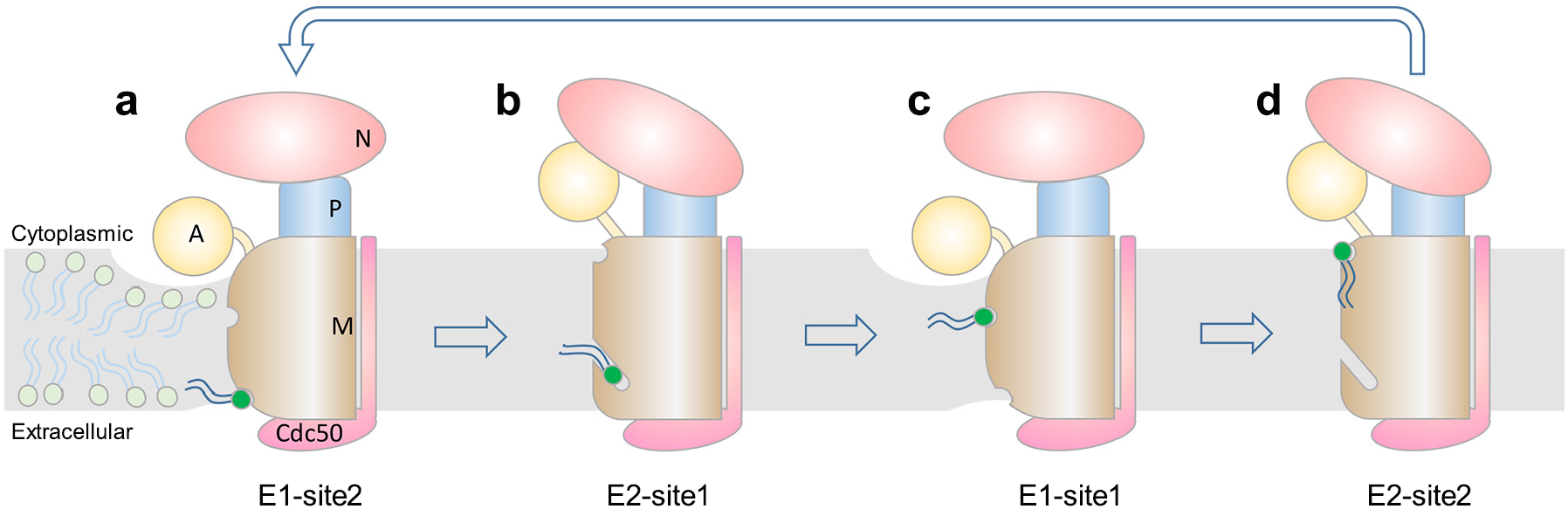
Model of phospholipid flipping by P4-ATPases. **a**, Cartoon drawing of E1-ATP. Domains are labeled and colored as in Fig 1**a**. The membrane is colored grey. The lipid molecules with light green heads are arranged to show the distortion of bilayers. The lipid molecule with the dark green head represents a substrate that is entering the transport pathway via E1-site2. **b**, Cartoon drawing of E2P. The lipid substrate is trapped in E2-site1. **c**, As in **a**, with the lipid bound at E1-site1. **d**, As in **b**, but the lipid substrate has been flipped to the cytosolic leaflet and waits at E2-site2 to be released.

Consistent with biochemical data^19^, the groove has high affinity to the lipid substrate in the E2P state as evident by the strong phospholipid density, whereas the groove shows weak lipid binding to facilitate lipid entry and exit in the E1-ATP state as indicated by the fragmented lipid density in the structure. Similarly, a hydrophilic membrane-traversing groove is also present in the TMEM16F scramblase^36^.

The “hydrophobic gate” model suggests that TMs 1 and 2 move away from TMs 3 and 4 during lipid transport^22^. Indeed, our structures show that TM1 and TM2 becomes flexible in E1-ATP. The key residue, I364 of the “hydrophobic gate” (I554 in ctDnf1), is at the interface between TMs 1, 2, and 4 (Fig. S11). Thus the mutations of the residue may disrupt the E1-E2 equilibrium and hamper lipid flipping as observed in the mutagenesis studies^22^. The “two-gate” model suggests that the flippases recognize the phospholipid substrates by interacting with the head groups. Residues other than the classical ion binding residues in ion-pumping P-type ATPases are involved in recognition^16,20,21,37^. Consistent with the model, the four distinct binding sites in our structures mainly interact with the lipid head groups. However, the sites do not seem to provide a discrimination mechanism for different phospholipid substrates. As shown in the phospholipid-dependent ATPase activity assays, ctDnf1 may have different substrate specificity from scDrs2 and hATP8A1. The amino acids that interact with the polar head group of the phospholipid substrate at E2-site1 are similar to those in the structures of scDrs2 and hATP8A1. The corresponding residues are Q549 and N550 of ctDnf12, S503 and N504 of scDrs2, and N352 and N353 of hATP8A1 (Fig. S1). The clamp residues of E2-site2 consist of both hydrophilic and hydrophobic residues, but are not conserved among P4-ATPases (Fig. S1). The serine residues at E1-site1 are highly conserved among P4-ATPases, and E1-site2 only provides a steric opening. Further studies on other intermediate states may provide clues on the substrate specificity.

In summary, the structures of ctDnf1-Cdc50 suggest that P4-ATPases have evolved a unique mechanism for lipid flipping. For ion-pumping P-type ATPases, most of TMs are involved in coordinating ion movements in the membranes. In contrast, TMs 3-10 of P4-ATPases are kept in a fixed conformation by Cdc50 during ATP hydrolysis. TMs 1 and 2 are the major regulators for lipid flipping. The distorted lipid bilayers and the groove bordered by TMs 2, 4, and 6 may be the key factors in controlling lipid flipping. It remains to be seen how the P4-ATPases select specific lipids to enter and exit the flipping pathway.

## Acknowledgments

We thank Dr. Xudong Wu and Dr. Qin Li for providing the yeast expression vectors, Dr. Stefan Schoebel for providing *Chaetomium thermophilum* cDNA, Dr. Ningning Li and Chengying Ma for helps on EM data analysis, Dr. Ning Gao and Dr. Ryan D. Baldridge for critical reading of the manuscript, the National Center for Protein Sciences at Peking University in Beijing, China, for assistance with protein purification, the Core Facilities at School of Life Sciences Peking University for assistance with negative staining EM, and the Electron Microscopy Laboratory of Peking University and the cryo-EM platform of Peking University for help with data collection. The computation was supported by the High-performance Computing Platform of Peking University. This work was supported by the National Natural Science Foundation of China (NSFC) (31870835 to L.L.), and the China Postdoctoral Science Foundation (2019M650327 to J.X.).

## Author Contributions

Y.H., J.X., and X.W. prepared the protein samples. Y.H. examined the ATPase activity. Y.H. and X.W. performed cryo-EM sample preparation and data collection. Y.H. and L.L. determined the structures. L.L. supervised the project. L.L., Y.H., J.X., and X.W. prepared the manuscript.

## Methods

### Protein expression and purification

The genes of ctDnf1 and ctCdc50 were cloned from the cDNA library of *Chaetomium thermophilum* (*var. thermophilum* strain: DSM1495, a gift from Dr. Stefan Schoebel). Superfolder green fluorescence protein (sfGFP)^38^, a Twin-Strep tag and a 3C protease cleavage site were fused to the N-terminus of ctDnf1. sfGFP, a His_9_ tag, and a 3C protease cleavage site were fused to the N-terminus of ctCdc50. The expression plasmids pRS426-sfGFP-twinStrep-3C-CtDnf1 and pRS424-sfGFP-His_9_-3C-CtCdc50 were co-transformed into *S. cerevisiae* strain BJ5465 using the LiAc/SS carrier DNA/PEG method^39^. Yeast cells were cultured in synthetic drop-out medium supplemented with 2% raffinose at 30 °C for about 24h to reach an optical density (OD_600_) of about 5. The culture was induced by the addition of 2% galactose and continued for 20 h at 25°C. The cells were harvested and stored at −80 °C until use.

The cells were suspended in the membrane extraction buffer (20 mM Tris-HCl pH 7.4, 150 mM NaCl, 5mM MgCl_2_,1mM DTT, and protease inhibitor cocktails) and lysed by high pressure homogenization. The crude lysate was clarified by centrifugation (20,000×g, 25 min, 4°C). The membrane fraction was pelleted by ultracentrifugation (200,000×g, 1 h, 4°C) and washed once with the membrane extraction buffer. The membrane pellets were solubilized in 2% lauryl maltose neopentyl glycol (LMNG, Anatrace) in the membrane solubilization buffer (20 mM Tris-HCl pH 7.4, 150 mM NaCl, 5mM MgCl_2_,1mM DTT, 10% glycerol, and protease inhibitor cocktails). After incubation at 4 °C for 1 h, the solution was clarified by ultracentrifugation (200,000×g, 1 h, 4°C). The supernatant was mixed with avidin (Sigma) and loaded onto a column pre-packed with StrepTactin resin (IBA Lifesciences). The eluents were concentrated and incubated with 3C protease at 4°C overnight. The protein solution was then loaded onto a Superdex 200 10/300 column (GE Healthcare). The peak fractions were pooled and concentrated (Fig. S2). The purified protein was either reconstituted into nanodiscs or flash-frozen in liquid nitrogen and stored at −80 °C.

The purified protein was mixed with MSP1D1^40^ and yeast polar lipids (Avanti Lipids, 40 mg/ml dissolved in 1% DDM) at a molar ratio of 1:2:25. Bio-beads SM2 (Bio-Rad) were then added to the mixture and incubated at 4 °C overnight to remove detergents. The complex was further purified by size-exclusion chromatography on a Superdex 200 10/300 column. The peak fraction had a protein concentration of 1.0 mg/ml (Fig. S2). It was immediately used for cryo-EM sample preparation without concentrating.

### Cryo-EM sample preparation and data collection

The freshly prepared samples were incubated with 1mM BeF_3_ or 1mM AMPPCP on ice for 30min before vitrification. The cryo-grid preparation was performed at 4 °C and 100% humidity in an FEI Vitrobot Mark IV. 4 µl sample was applied to each freshly glow-discharged grid (Quantifoil, R1.2/1.3). The grids were then plunge-frozen in liquid ethane. The cryo-grids were screened with a 200 kV FEI Talos Arctica microscope equipped with a FEI Ceta camera. The data were collected on a 300 kV FEI Titan Krios TEM with a K2 summit camera and GIF Quantum energy filter (Gatan). The images were collected at a magnification of 130,000× with a calibrated pixel size of 1.055 Å. The dose rate was set at 8 e^−^/s/pixel and the exposure time was 8 s, corresponding to a total dose of 57.5 e^−^/Å^2^. Movie stacks (32 frames each) were recorded with the software SerialEM^41^ under low-dose conditions with defocuses ranging from −1 to −2 μm.

### Image processing

The movie stacks were subject to motion correction and electron-dose weighting by using MotionCor2^42^ (Fig. S3a, S4a). The program Gctf^43^ was used to estimate the contrast transfer function (CTF) parameters. Images of high quality were selected for further image processing on the basis of the CTF power spectra of the corrected images. The following calculations are performed with RELION3.0^44^. Particles of high quality were selected according to 2D classification (Fig. S3b, S4b) and 3D classification results. The selected particles were subject to several rounds of CTF refinement and polishing. After mask-based post-processing, the final maps had resolutions of 3.40 Å and 3.48 Å for the AMPPCP and BeF_3_^-^ samples, respectively (Fig. S3, S4). All the resolution estimations were based on gold-standard Fourier Shell Correlation (FSC) 0.143 criteria.

The model for the E2P (BeF_3_^-^) structure was built manually in Coot^45^, with the guidance of the scDrs2 structures. The model was refined in real space using Phenix^46^. For the model building of E1-ATP (AMPPCP), the E2P model was fit in the E1-ATP density map. Each domain is subject to rigid body refinement. Due to the local resolution limits, the A and N domains were not refined further. The rest parts of the E1-ATP model were refined in real space with Phenix. Model validation was done with MolProbity^47^.

### ATPase activity assay

The ATPase activity assays were carried out by using BIOMOL® Green (Enzo) to measure the free phosphate concentrations. The reaction solutions consisted of 0.05mg/ml protein, 0.01% LMNG, 0.02% C_12_E_9_ (Anatrace), 150 mM NaCl, 20 mM HEPES-NaOH pH 7.5, 5mM MgCl_2_, 1mM DTT, 2.5mM ATP, and lipids at the indicated concentrations. The reactions were carried out at 30 °C for 20 min, and then immediately diluted 10 times for color development. 100 µl reagent was added to 50 µl sample and the mixture was incubated at room temperature for 20 min. The absorbance at 650 nm was measured in a microplate reader (BioTek Cytation5). The phosphate concentration was determined by calibration with the phosphate standard (BML-KI102).

---

**Table S1.** Cryo-EM data collection, refinement and validation statistics.

|  | ctDnf1-Cdc50<br>with AMPPCP | ctDnf1-Cdc50<br>with BeF<sub>3</sub><sup>-</sup> |
| --- | --- | --- |
| **Data collection and processing** |  |  |
| Magnification | 130,000 | 130,000 |
| Voltage (kV) | 300 | 300 |
| Electron exposure (e<sup>-</sup>/Å<sup>2</sup>) | 57.5 | 57.5 |
| Defocus range (μm) | -1.0 to -2.0 | -1.0 to 2.0 |
| Pixel size (Å) | 1.055 | 1.055 |
| Symmetry imposed | C1 | C1 |
| Initial particle images (no.) | 2,397,258 | 2,820,251 |
| Final particle images (no.) | 272,912 | 249,694 |
| Map resolution (Å) | 3.40 | 3.48 |
| FSC threshold 0.143 |  |  |
| Map resolution range (Å) | 3.3-8.2 | 3.3-7.8 |
| **Refinement** |  |  |
| Model resolution (Å) | 3.4 | 3.5 |
| Map sharpening B factor (Å<sup>2</sup>) | -94 | -84 |
| **Model composition** |  |  |
| Non-hydrogen atoms | 11590 | 12406 |
| Protein residues | 1420 | 1514 |
| Ligands | 17 | 18 |
| **B factors (Å<sup>2</sup>)** |  |  |
| Protein | 41.2 | 50.0 |
| Ligand | 45.6 | 51.5 |
| **R.m.s. deviations** |  |  |
| Bond lengths (Å) | 0.012 | 0.010 |
| Bond angles (°) | 1.055 | 1.088 |
| **Validation** |  |  |
| MolProbity score | 2.16 | 2.35 |
| Clashscore | 15.5 | 18.1 |
| Poor rotamers (%) | 0.31 | 0.25 |
| **Ramachandran plot** |  |  |
| Favored (%) | 89.8% | 88.1% |
| Allowed (%) | 10.2% | 11.7% |
| Disallowed (%) | 0% | 0.2% |

**Fig. S1.**
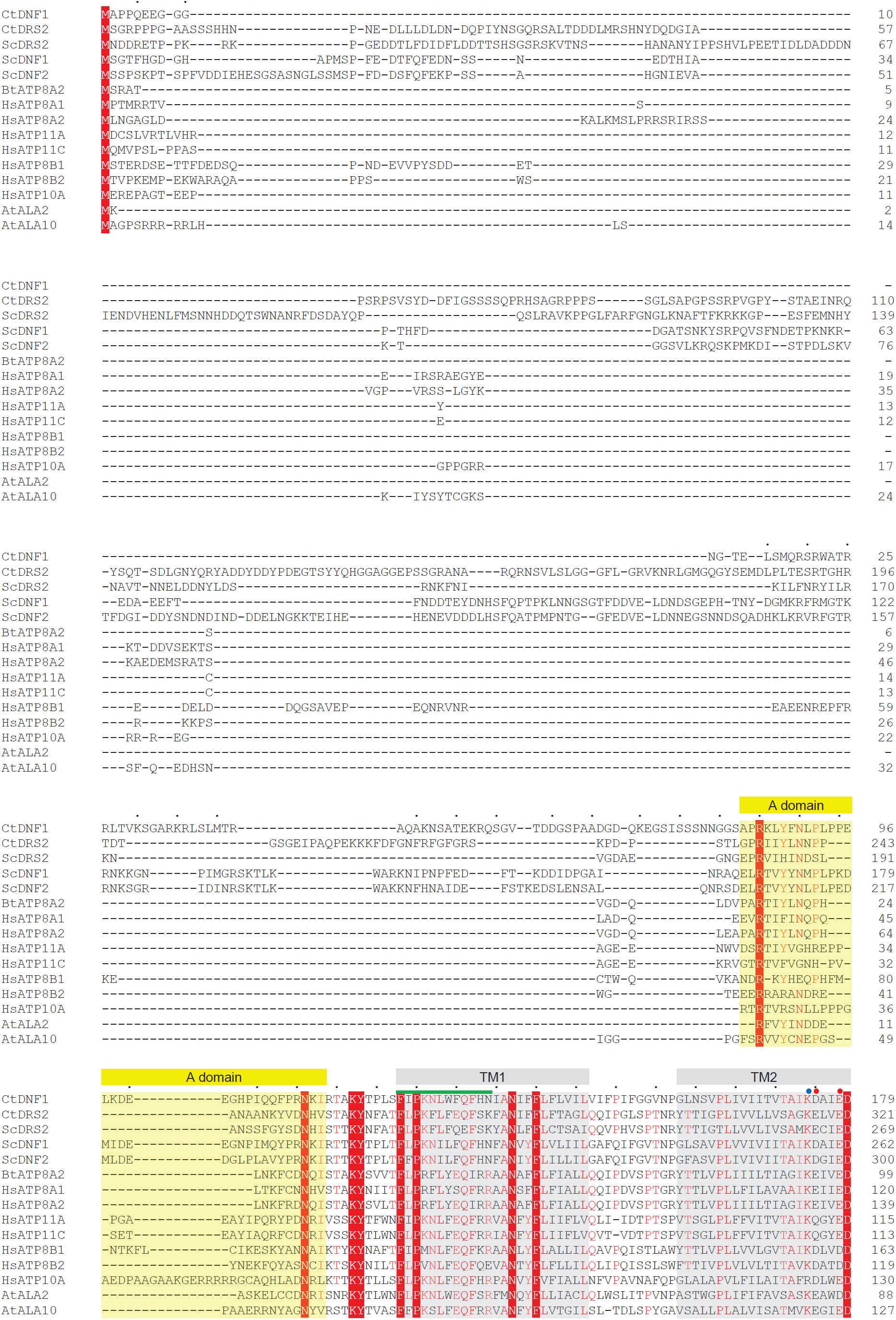
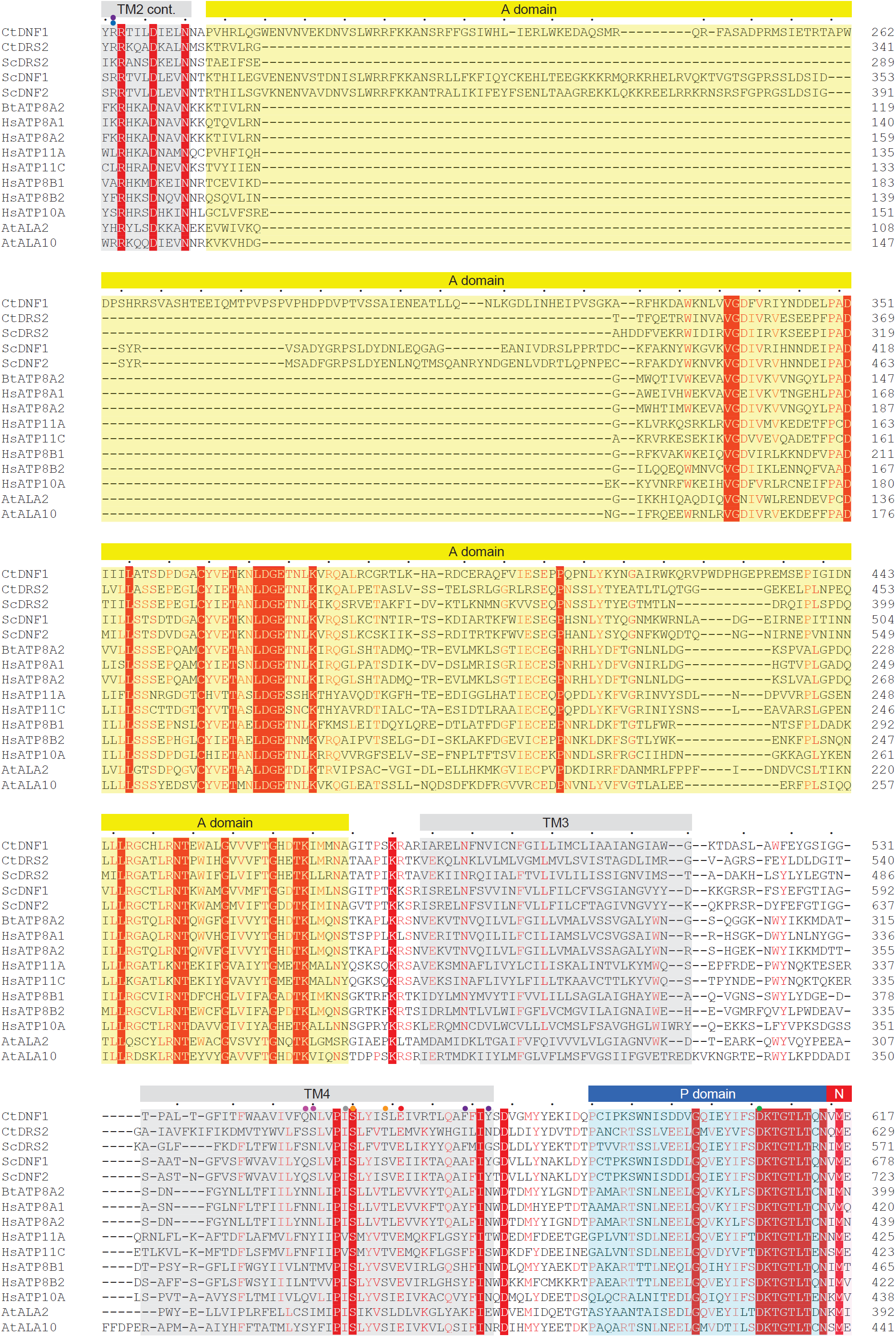
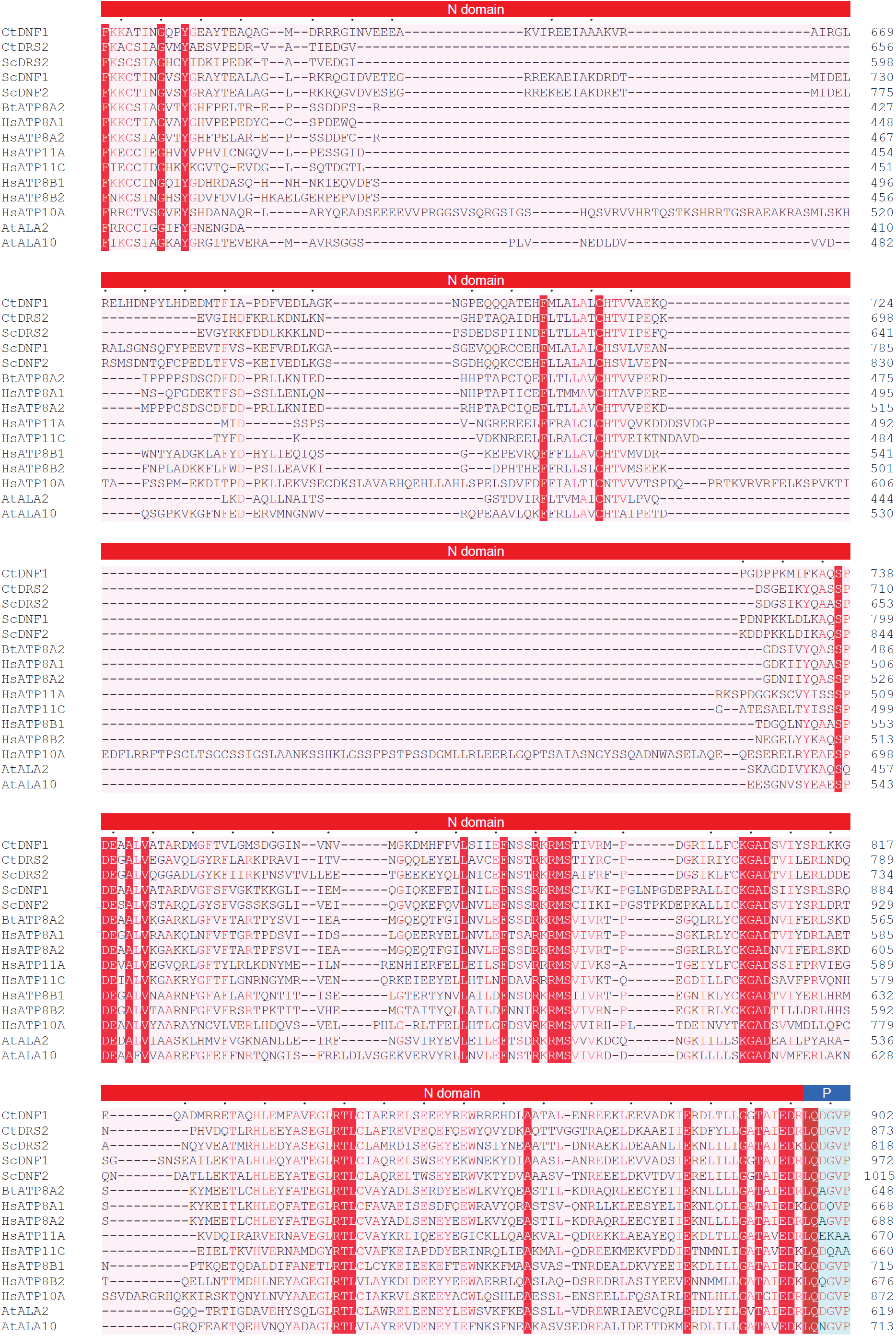
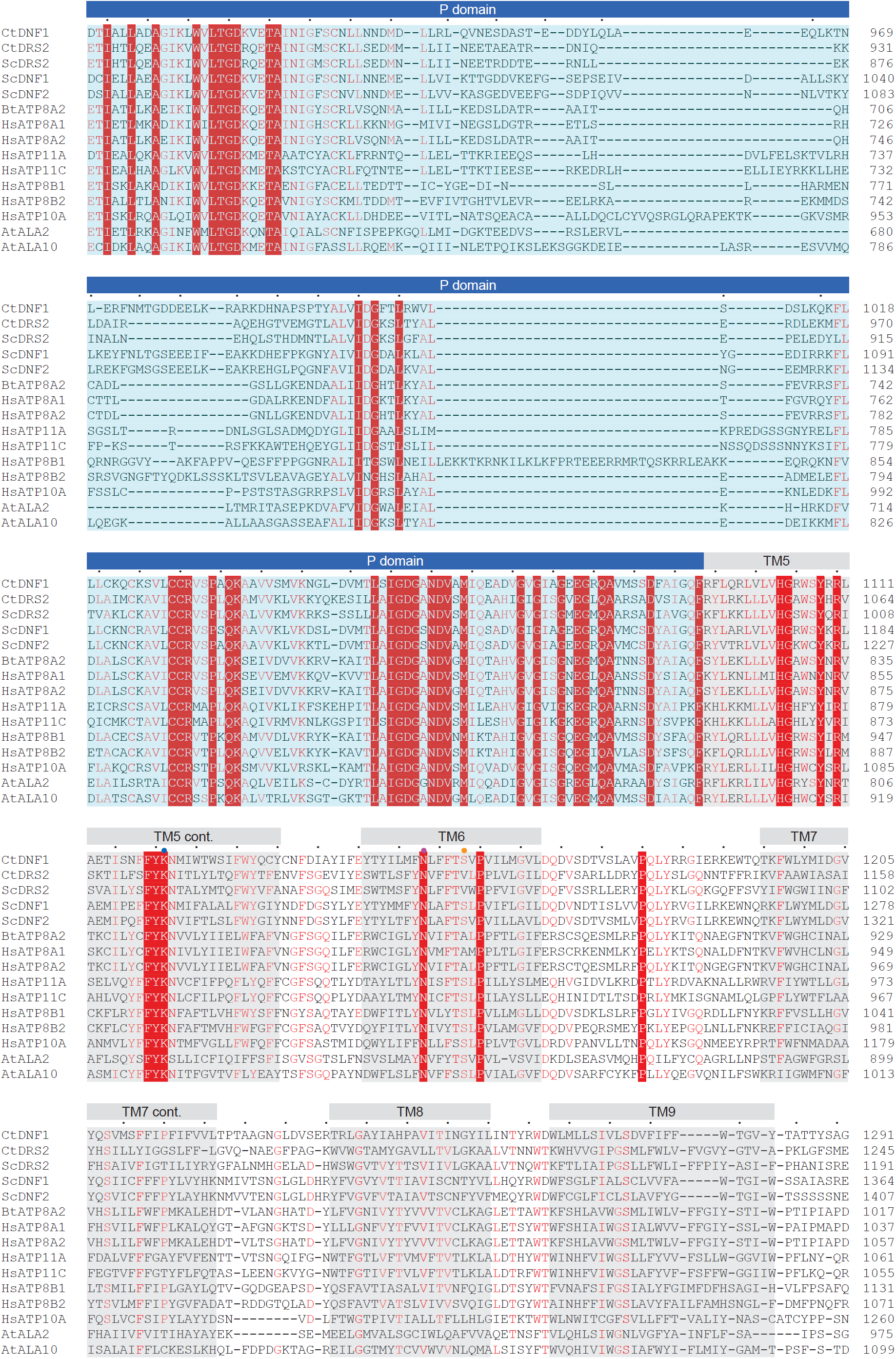
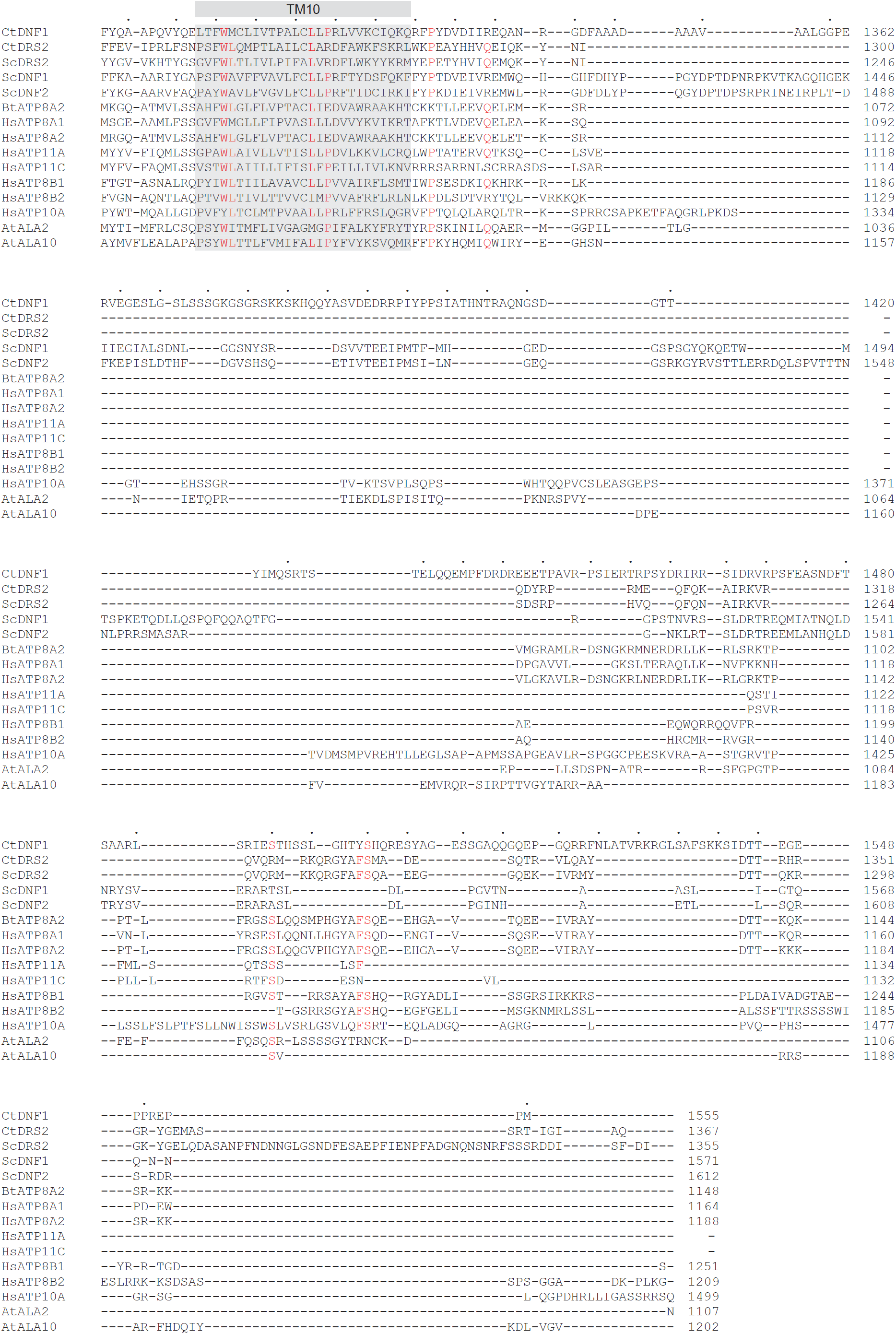
Sequence alignment of selected P4-ATPases. Sequence alignment of ctDnf1, ctDrs2 and other P4-ATPases in yeast, bovine, human, and A. thaliana, aligned by T-coffee^48^. The conserved domains and transmembrane helices of ctDnf1 are indicated above the sequences. The conserved residues are indicated in red letters. The amphipathic helix of TM1 is highlighted with a green bar above the alignment. The phosphorylation site of the P domain is highlighted with a green dot. The residues involved in the negatively charged patch are highlighted with red dots. The residues contribute to the positive charge of the groove are highlighted with blue dots. The residues involved in E1-site1 are highlighted with orange dots. The residues involved in E2-site1 are highlighted with magenta dots. The residues involved in E1-site2 are highlighted with purple dots. The key isoleucine residue in the hydrophobic gate model is highlighted with a grey dot. Ct, *Chaetomium thermophilum*; Sc, S*accharomyces cerevisiae*; Bt, *Bos Taurus*; Hs, *Homo sapiens*; At, *Arabidopsis thaliana*. Uniprot accession numbers: ScDRS2, P39524; ScDNF1, P32660; ScDNF2, Q12675; BtATP8A2, C7EXK4; HsATP8A1, Q9Y2Q0; HsATP8A2, Q9NTI2; HsATP11A, P98196; HsATP11C, Q8NB49; HsATP8B1, O43520; HsATP8B2, P98198; HsATP10A, O60312; AtALA2, P98205; AtALA10, Q9LI83.

**Fig. S2.**
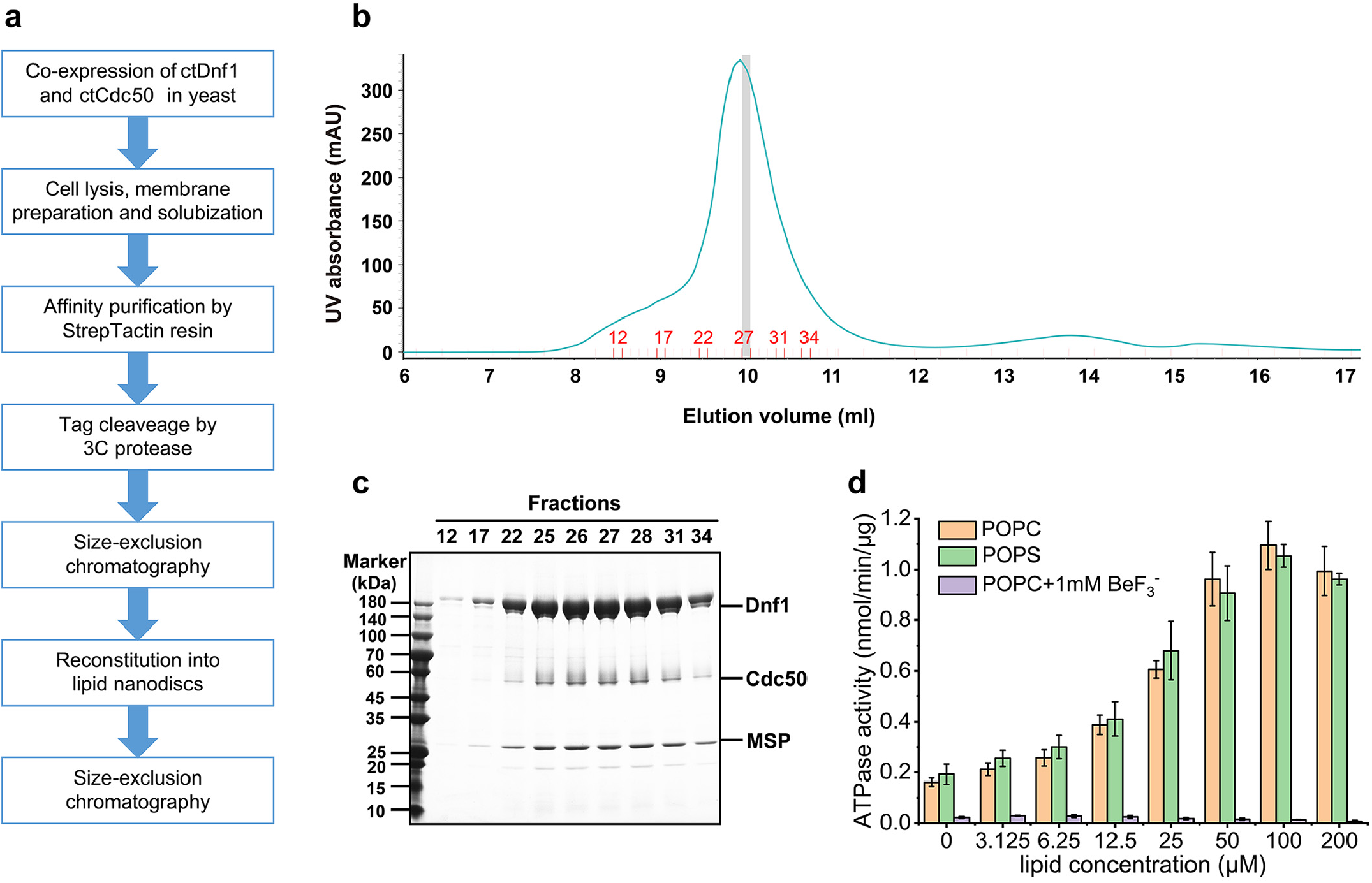
Purification and characterization of the ctDnf1-Cdc50 complex. **a**, Flow chart of ctDnf1-Cdc50 purification. **b**, Size-exclusion chromatography profile of the protein complex reconstituted into nanodiscs. The gray-shaded area (Fraction 27) was used for cryo-EM analysis. **c**, SDS-PAGE analysis of fractions from the SEC purification in **b. d**, ATPase activity of ctDnf1-Cdc50 complex stimulated by phospholipids. Data points represent the mean ± SEM of at least three experiments. POPC, 1-palmitoyl-2-oleoyl-sn-glycero-3-phosphocholine. POPS, 1-palmitoyl-2-oleoyl-sn-glycero-3-phospho-L-serine.

**Fig. S3.**
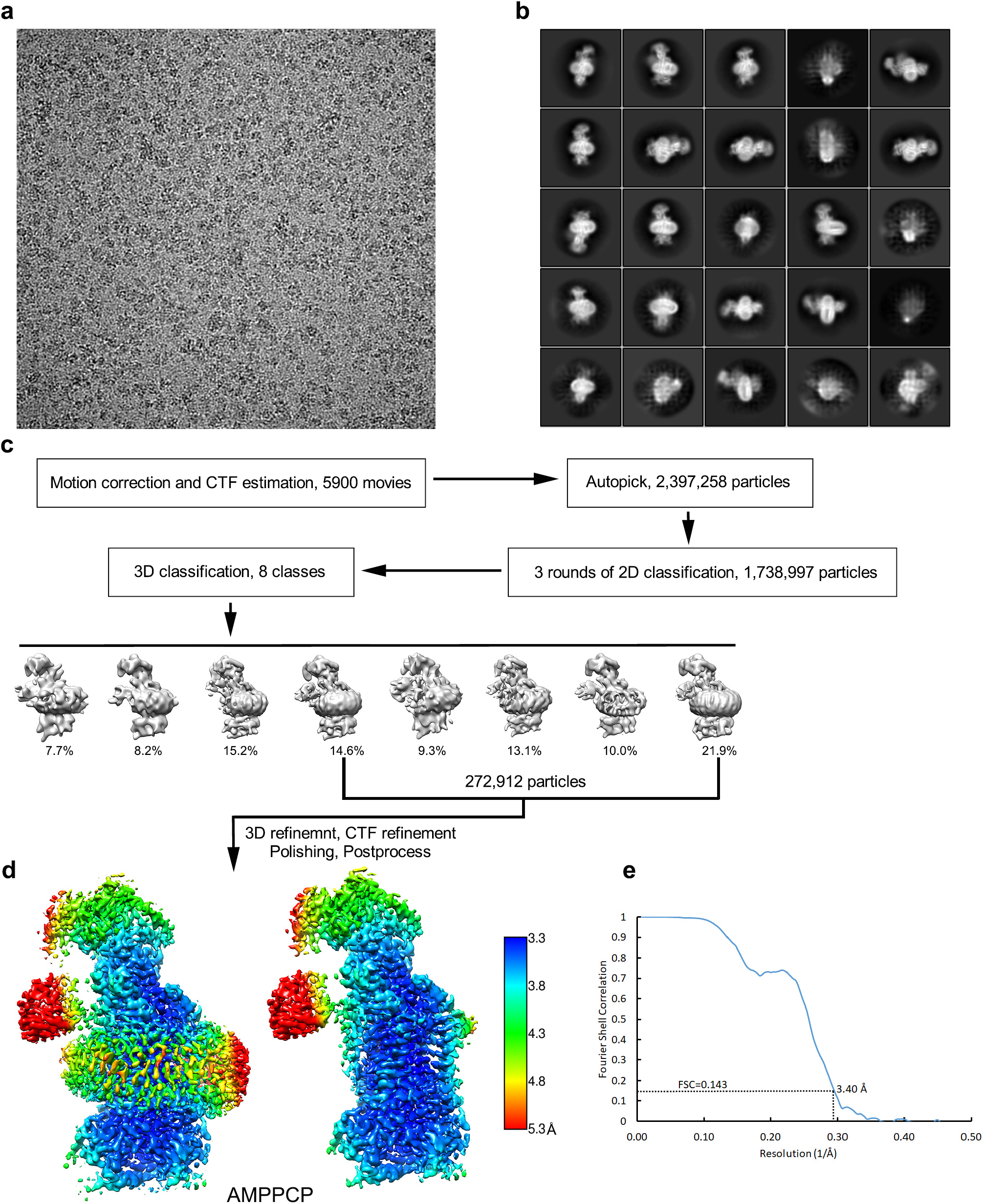
Cryo-EM single particle analysis of ctDnf1-Cdc50 with AMPPCP. **a**, Representative image after motion correction. **b**, Representative results of 2D classification. **c**, Workflow of the single particle analysis. **d**, Local resolution map of the final sharpened map, shown with (left) and without (right) the nanodisc. **e**, Fourier shell correlation (FSC) curve with estimated resolution according to the gold standard.

**Fig. S4.**
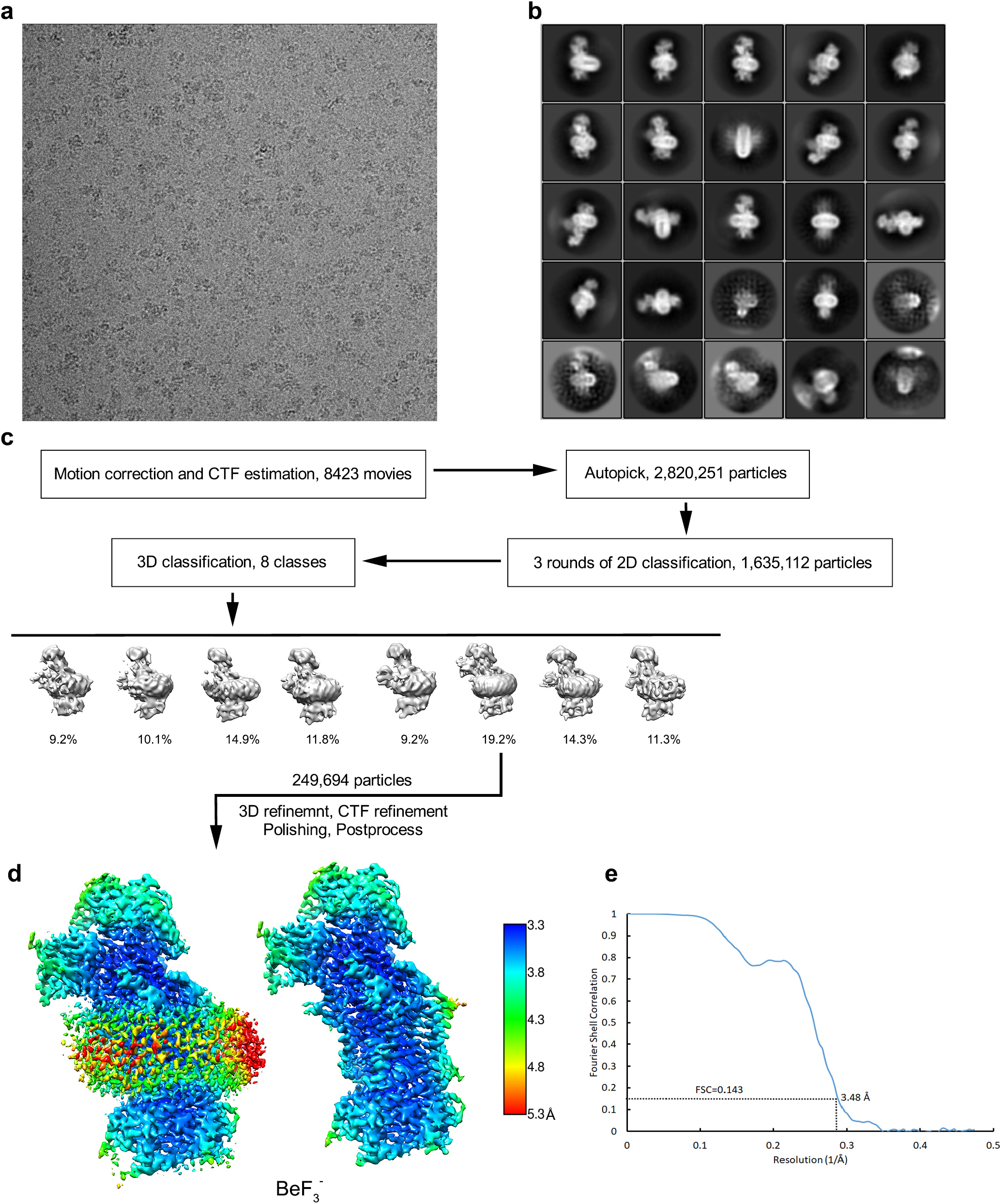
Cryo-EM single particle analysis of ctDnf1-Cdc50 with BeF_3_^-^. **a**, Representative image after motion correction. **b**, Representative results of 2D classification. **c**, Workflow of the single particle analysis. **d**, Local resolution map of the final sharpened map shown with (left) and without (right) the nanodisc. **e**, Fourier shell correlation (FSC) curve with estimated resolution according to the gold standard.

**Fig. S5.**
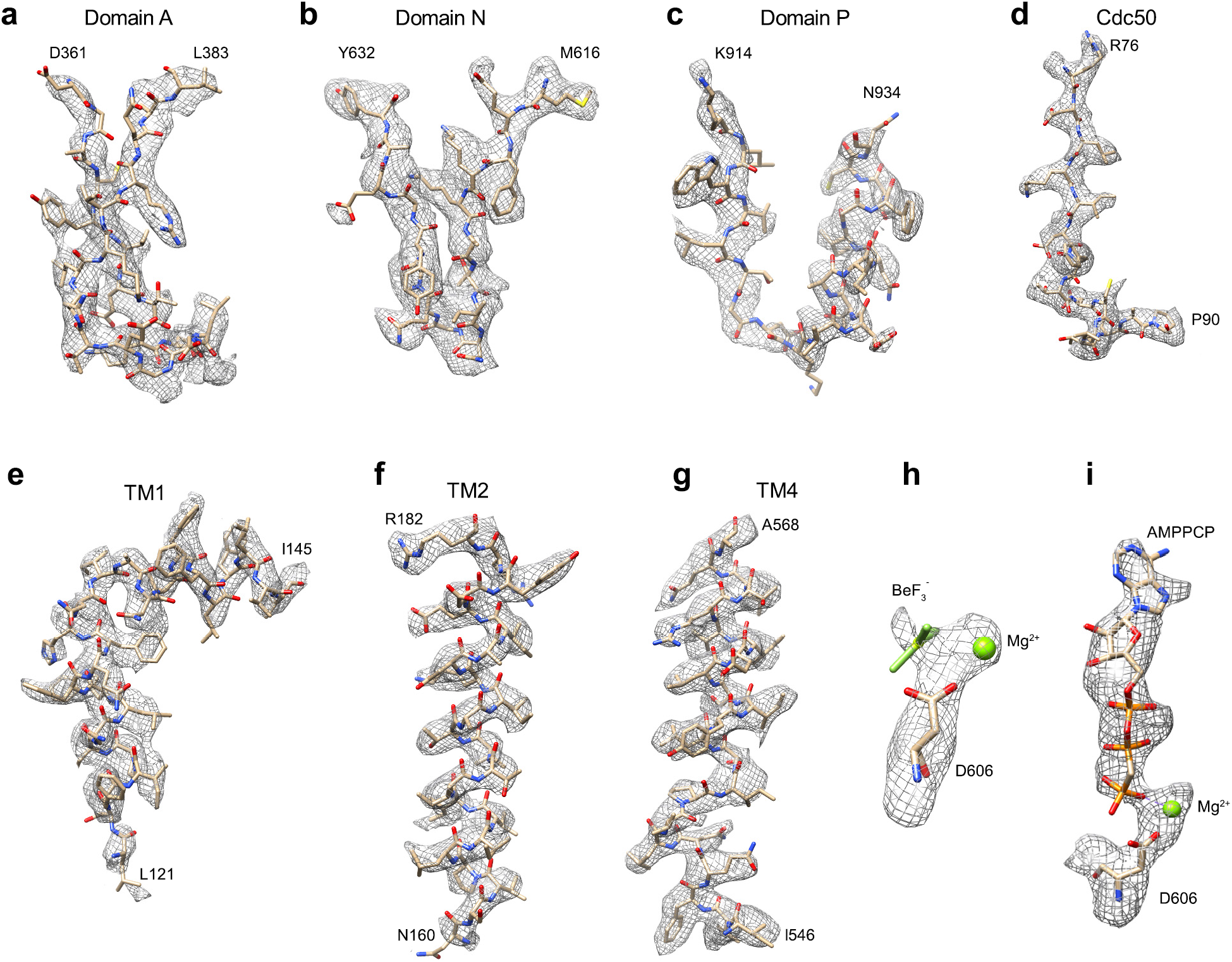
Examples of the fit of models into the density map. **a-g**, Density map and model in selected regions of each domain and TMs. Residues at the beginning and end of each polypeptide segment are indicated. **h**, Density map and model of BeF_3_^-^, Mg^2+^, and D606 in the E1-ATP structure. **i**, Density map and model of AMPPCP, Mg^2+^, and D606 in the E1-ATP structure.

**Fig. S6.**
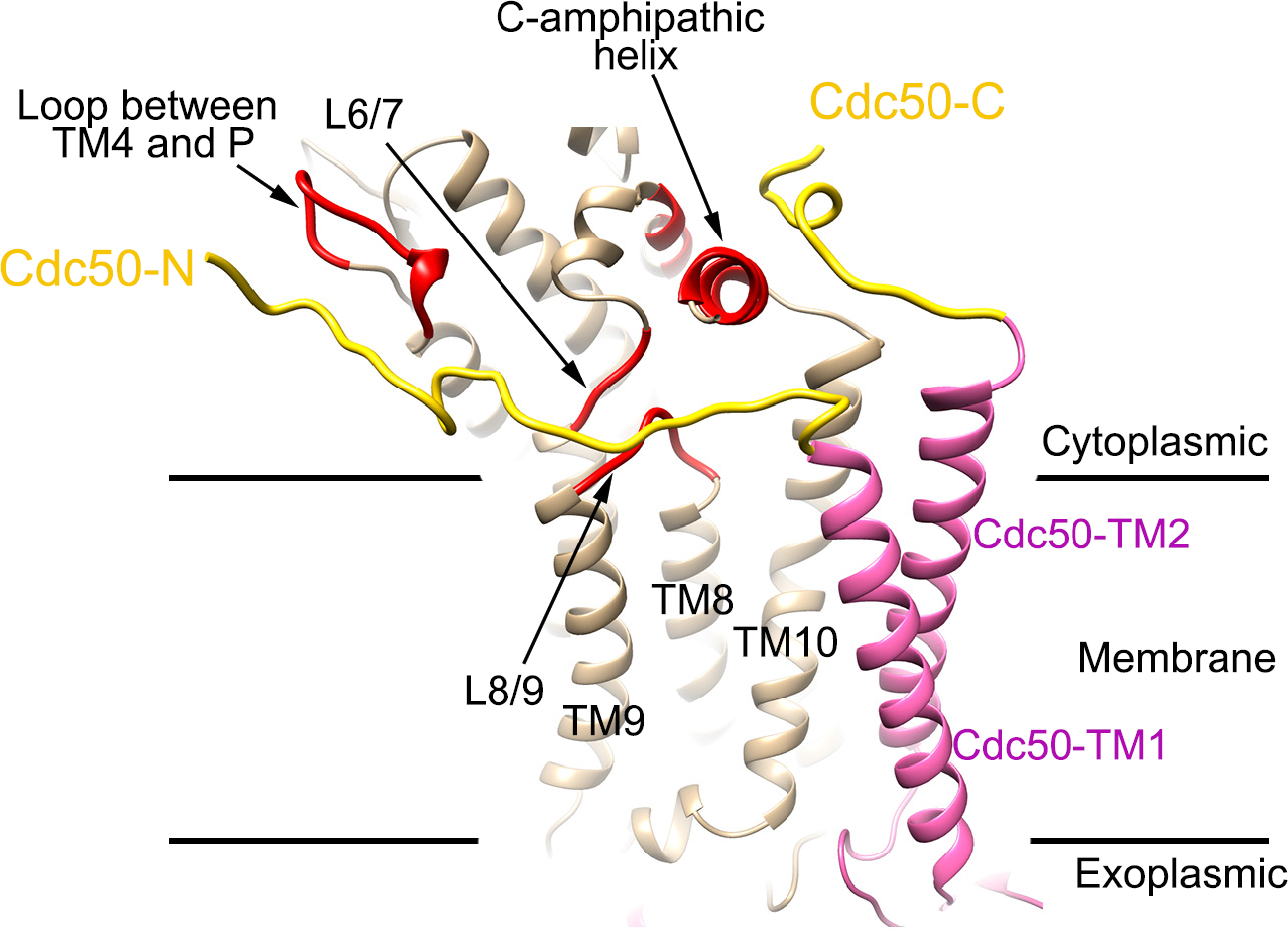
Interaction of the two terminal segments of ctCdc50 with ctDnf1 on the cytoplasmic side. The N-terminal and C-terminal segments of ctCdc50 are colored yellow. The rest of ctCdc50 is pink. The ctDnf1 fragments that interact with ctCdc50 are colored red. The rest of ctDnf1 is tan. Interacting segments and TMs are labeled.

**Fig. S7.**
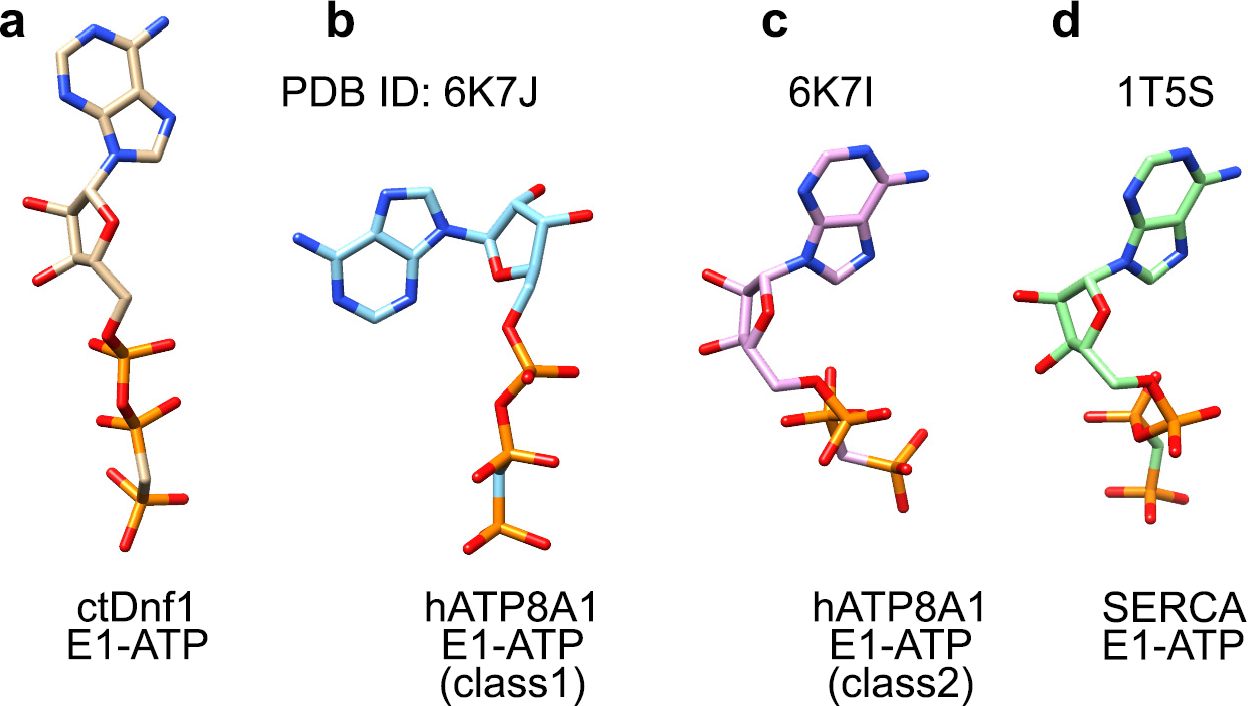
Comparison of AMPPCP in different E1-ATP structures. **a-d**, AMPPCP conformations from different P-type ATPases are shown as sticks. The protein structures from which AMPPCP are extracted are labeled.

**Fig. S8.**
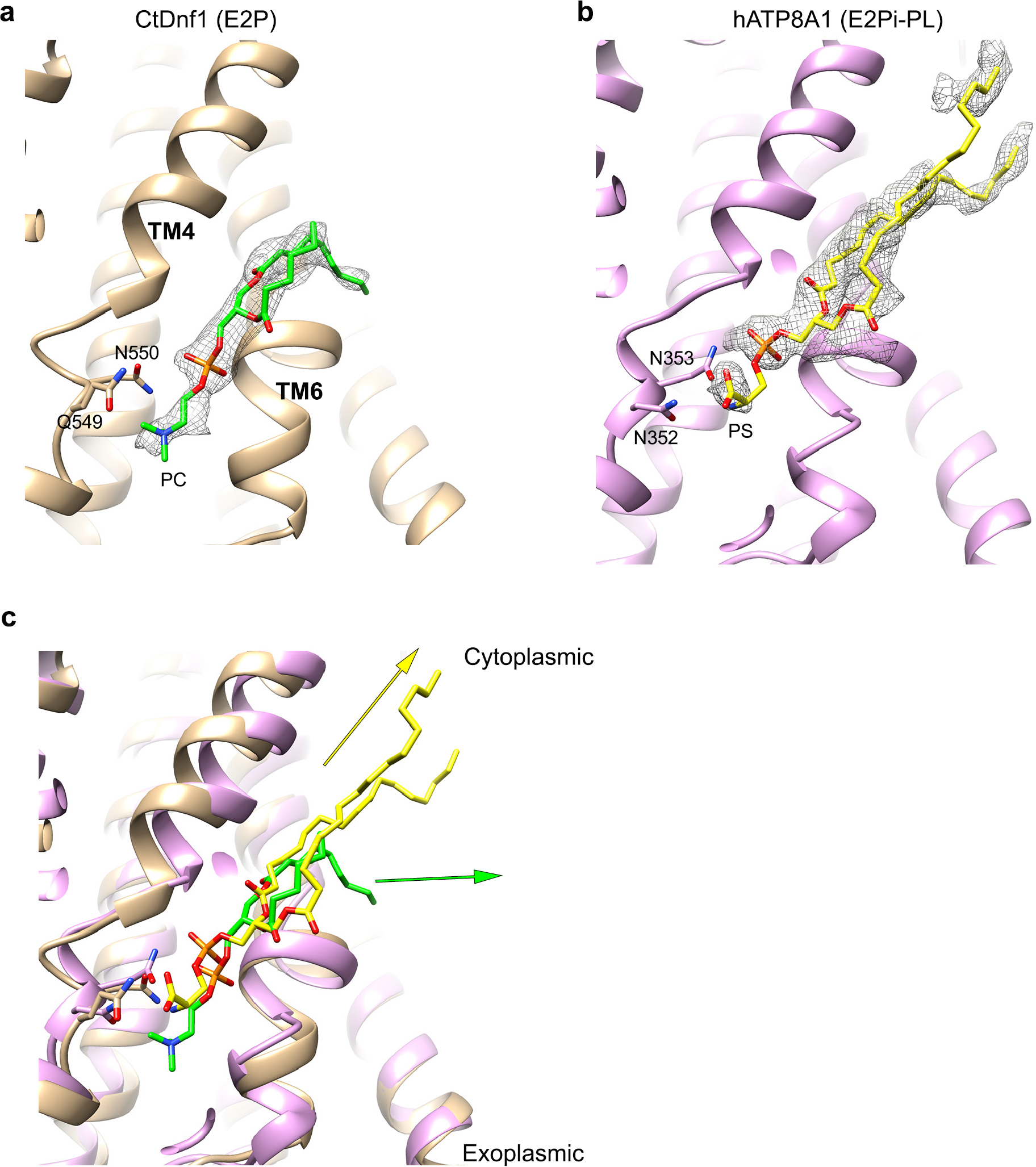
Comparison of phospholipid binding at E2-site1 in ctDnf1 and hATP8A1. **a**, Lipid binding at E2-site1 of ctDnf1. The protein is shown as tan ribbon representation. The density of the lipid is shown as a grey mesh. The lipid (green) and its interacting residues (tan) are shown as sticks. **b**, Lipid binding in the E2Pi-PL structure of hATP8A1 (PDB ID: 6K7M, EMDB number: 9941). The protein is colored purple. The lipid is yellow. **c**, Superimposition of the two lipid binding sites. The yellow arrow and green arrow indicate the extension directions of the lipid acyl chains in hATP8A1 and ctDnf1, respectively.

**Fig. S9.**
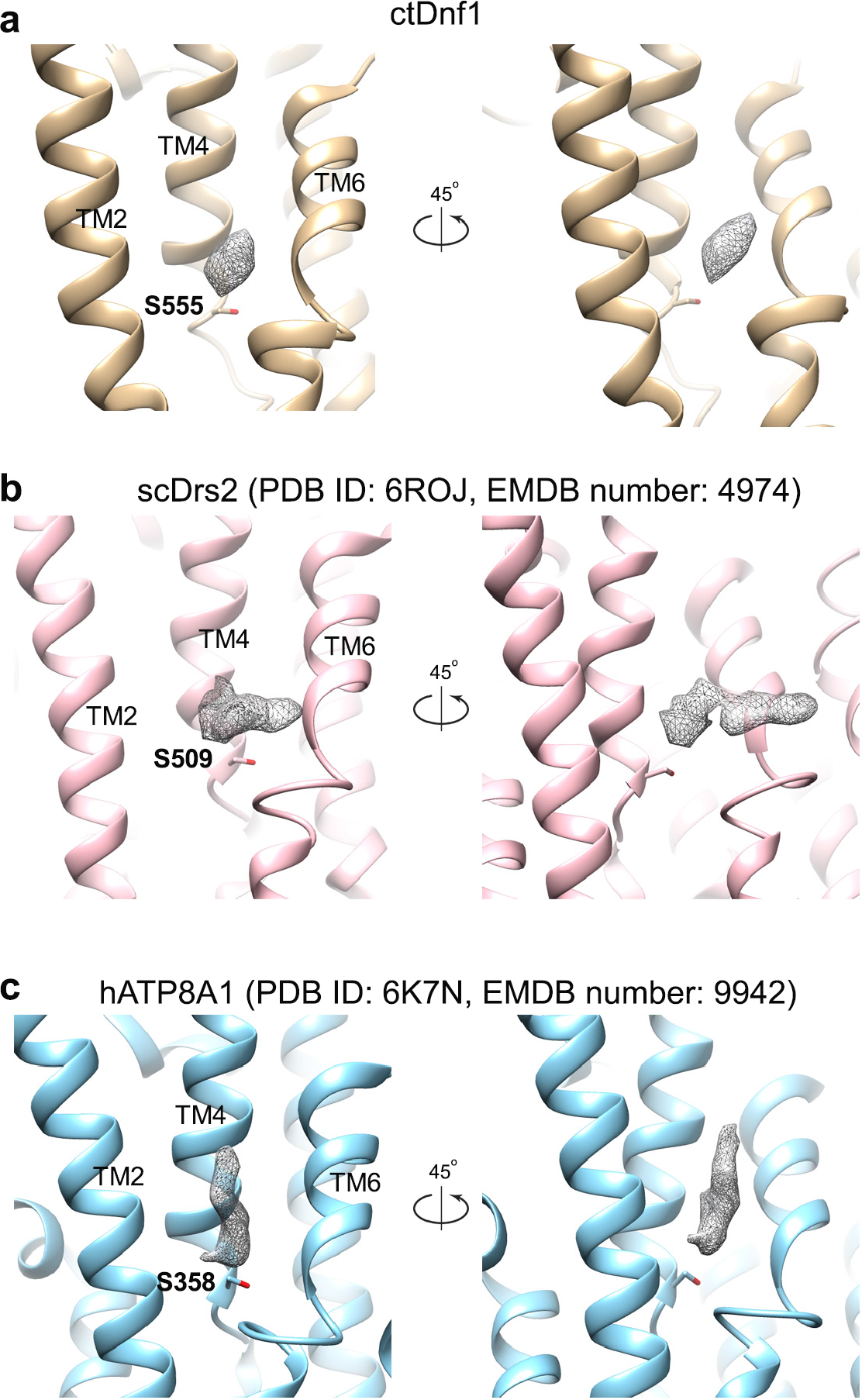
Comparison of E1-site1 among P4-ATPase structures. **a**, E1-site1 in ctDnf1. The density of the possible phospholipid substrate is shown as a grey mesh. The conserved serine residue is labeled. **b-c**, same as in **a**, except showing scDrs2 and hATP8A1, respectively.

**Fig. S10.**
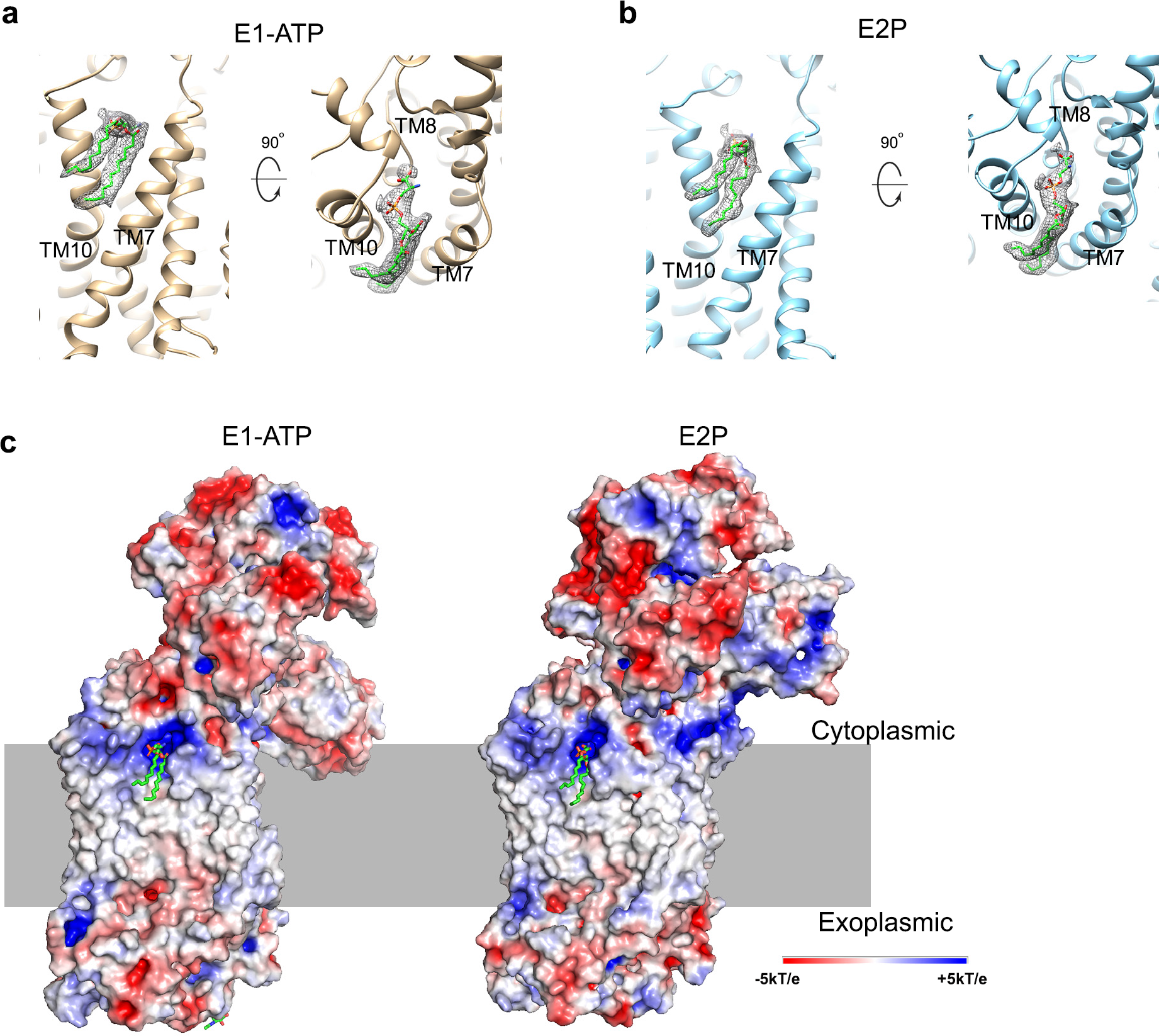
A common lipid binding site in E1-ATP and E2P. **a-b**, Lipid binding site in E1-ATP (**a**) and E2P (**b**). The density of the lipid is shown as grey meshes. The lipid molecules are shown as sticks. **c**, Electrostatic potential surfaces of E1-ATP (left) and E2P (right), showing the lipid binding environment. The lipid molecules are shown as sticks. The surfaces showing here are on the opposite side of the surfaces showing in Fig. 4b and d.

**Fig. S11.**
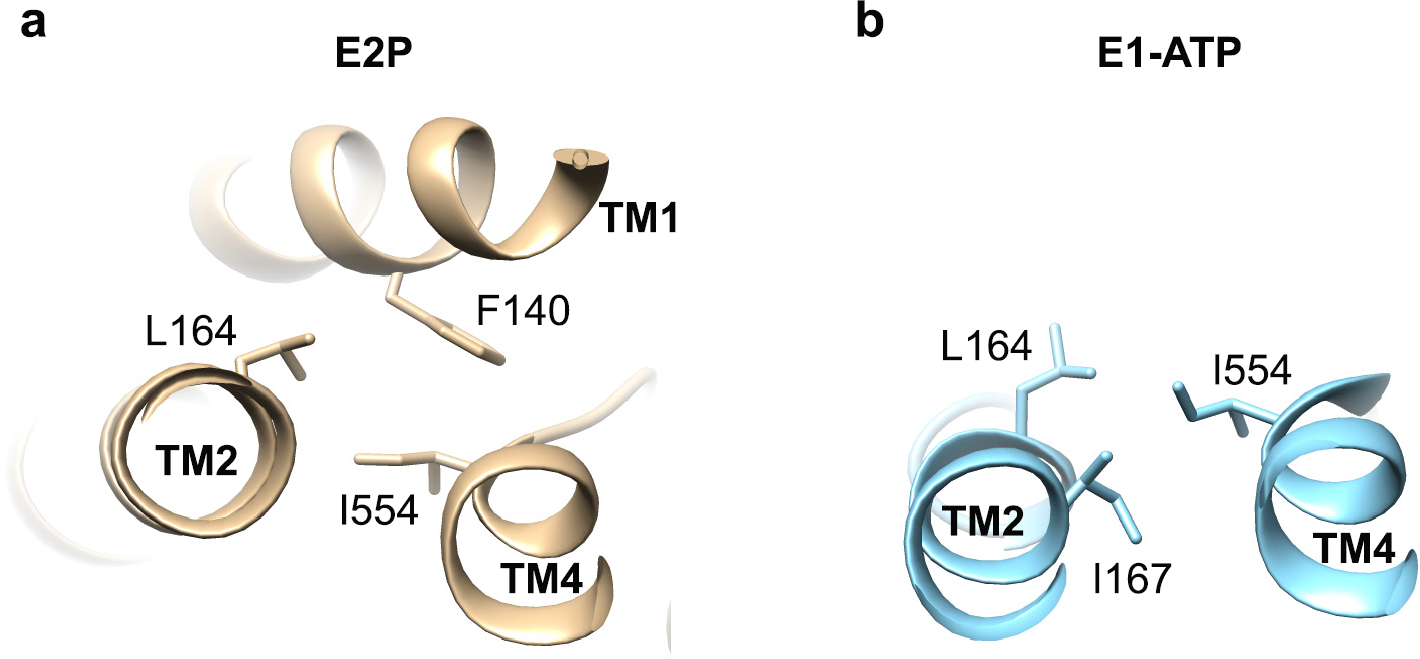
The “hydrophobic gate” residue I554 in cfDnf1. **a**, Top view of the I554 and its interacting residues in E2P. The interacting residues are shown as sticks. **b**, Top view of I554 in E1-ATP. TM4 is in the same orientation as it is in a.

